# Inhibition of striatal SEZ6 by miR-3594-5p is a drug-specific marker for late-stage heroin intake escalation

**DOI:** 10.1101/2021.07.26.453355

**Authors:** Magalie Lenoir, Isabella Bondi, Loïc Clemenceau, Isabelle Nondier, Margaux Ballé, Sébastien Jacques, Angéline Duché, Corinne Canestrelli, Séverine Martin-Lannerée, Sophie Mouillet-Richard, Jenny M. Gunnersen, Serge H. Ahmed, Nicolas Marie, Florence Noble

## Abstract

Escalation of drug use is a hallmark stage in the transition to addiction and uncovering its underlying brain molecular mechanisms constitutes a considerable challenge. Here, we report in rats with extended access to heroin for self-administration that miR-3594-5p was upregulated in the dorsal striatum at late, but not early, stages during escalation when excessive heroin intake plateaued. Striatal miR-3594-5p bound directly to the 3’UTR region of *Sez6* transcript and inhibited its expression, thereby decreasing the mature form of the translated SEZ6 protein. This miR-3594-5p/*Sez6* interaction was specific to heroin, as it was not observed with cocaine, and correlated with the severity of heroin intake escalation. Our findings reveal that miRNA alterations during escalation of drug self-administration are spatially and temporally regulated and drug-specific.

## INTRODUCTION

Drug addiction is mainly characterized by continued excessive drug use despite harmful consequences. Discovering brain molecular mechanisms that underlie the transition to addiction is important because it is a first step toward the definition of new therapeutic targets. Over the past two decades, several animal models of the transition from recreational drug use to addiction have been developed to facilitate the research of its neurobiological underpinnings^1^. In one of the most studied models, rats are given limited versus extended access to the drug for self-administration. Only rats with extended access (e.g., 6 h/day, Long Access or LgA rats) escalate their drug consumption over days while those with limited access (e.g., 1 h/day, Short Access or ShA rats) show a stable drug intake. LgA rats, but not ShA rats, also develop several other behavioral changes that resemble some of the behavioral symptoms of addiction^2^. They work harder and take more risk to seek and to consume the drug, they show difficulty of abstaining from drug seeking, they are more sensitive to drug and stress-primed craving behavior, and they also display some neurocognitive deficits.

Transition from recreational drug use to addiction is associated with persistent alterations in brain synaptic plasticity, including drug-induced alterations in long-term potentiation (LTP) and long-term depression (LTD) ^3, 45^. These long-lasting alterations have been observed in the nucleus accumbens (NAc)^4^, a ventral striatal area that is part of the brain reward system, as well as in the dorsal striatum (DS)^6^, which is part of the habit system^7–9^. Long-lasting structural and functional modifications underlying addiction-related synaptic plasticity require epigenetic mechanisms, which include regulatory non-coding microRNAs (miRNA), a class of molecular regulators of gene expression^10, 11^. miRNAs regulate gene expression by binding to complementary sequence in the 3’ untranslated region of target (3’UTR) mRNA transcripts to facilitate their degradation and/or inhibit their translation. miRNAs are of particular interest because they modulate many molecular and cellular adaptations involved in drug addiction^12–17^. In a seminal study, the expression of some miRNAs (i.e., miR-212 and miR-132) was shown to be enhanced only in the DS of LgA rats which had developed cocaine intake escalation following extended access, but not in ShA rats^18^. Interestingly, miR-212 overexpression was a molecular counteradaptation, via its action on CREB and MeCP2/BDNF signaling^18, 19^, to escalated levels of cocaine intake, thereby supporting its critical role as a potential neuroprotective mechanism in cocaine addiction.

Unlike cocaine, the role of miRNAs in opioid addiction and escalation has been relatively little studied^20–27^. Notably, we do not know whether heroin intake escalation after extended access is associated with the same or different miRNA alterations as those found previously with cocaine. Here we identified miRNA alterations in NAc and DS specific to LgA rats after extended access to heroin^1, 28–30^ and compared them directly to those induced by cocaine in the same experimental conditions. This research allowed us to discover for the first time the involvement of a new target gene, *Sez6,* specific to heroin intake escalation.

## RESULTS

### Extended access to heroin increases miR-3594-5p expression in the dorsal striatum

In Experiment 1 (Fig. 1a and Extended Data Fig. 2a), to identify transcriptional impairments implicated in the transition to heroin addiction, we first profiled microRNA expression in DS (Fig. 1 and Extended Data Fig. 1) and NAc (Extended Data Fig. 2) following 20 days of differential access to heroin for self-administration using Affymetrix Genechip miRNA 4.0 arrays. As expected^28, 29^, Long-Access (LgA) rats gradually increased their heroin intake from 5 to 19 infusions during the first hour of the long access sessions of heroin self-administration, whereas Short-Access (ShA) rats developed a stable and limited consumption (two-way repeated measures ANOVA: Group x Session: F_19, 152_ = 7.59, P<0.0001,Fig. 1b). In LgA rats, heroin injections during the last 5 hours also increased over time from 3 to 23 infusions at doses of 60 µg (F_3.073,12.29_=14.73, P=0.0002; Extended Data Fig. 1a). Rats in the LgA-Yoked group passively received the heroin infusions that their matched LgA rats obtained by self-administration (see Methods for experimental details). In contrast to ShA and LgA rats, these rats did not learn the operant behavior even after the 20 self-administration sessions, as indicated by the lack of discrimination between active and inactive levers (Extended Data Fig. 1b, c; see Supplementary Table 3 for Statistical details).

**Fig. 1.**
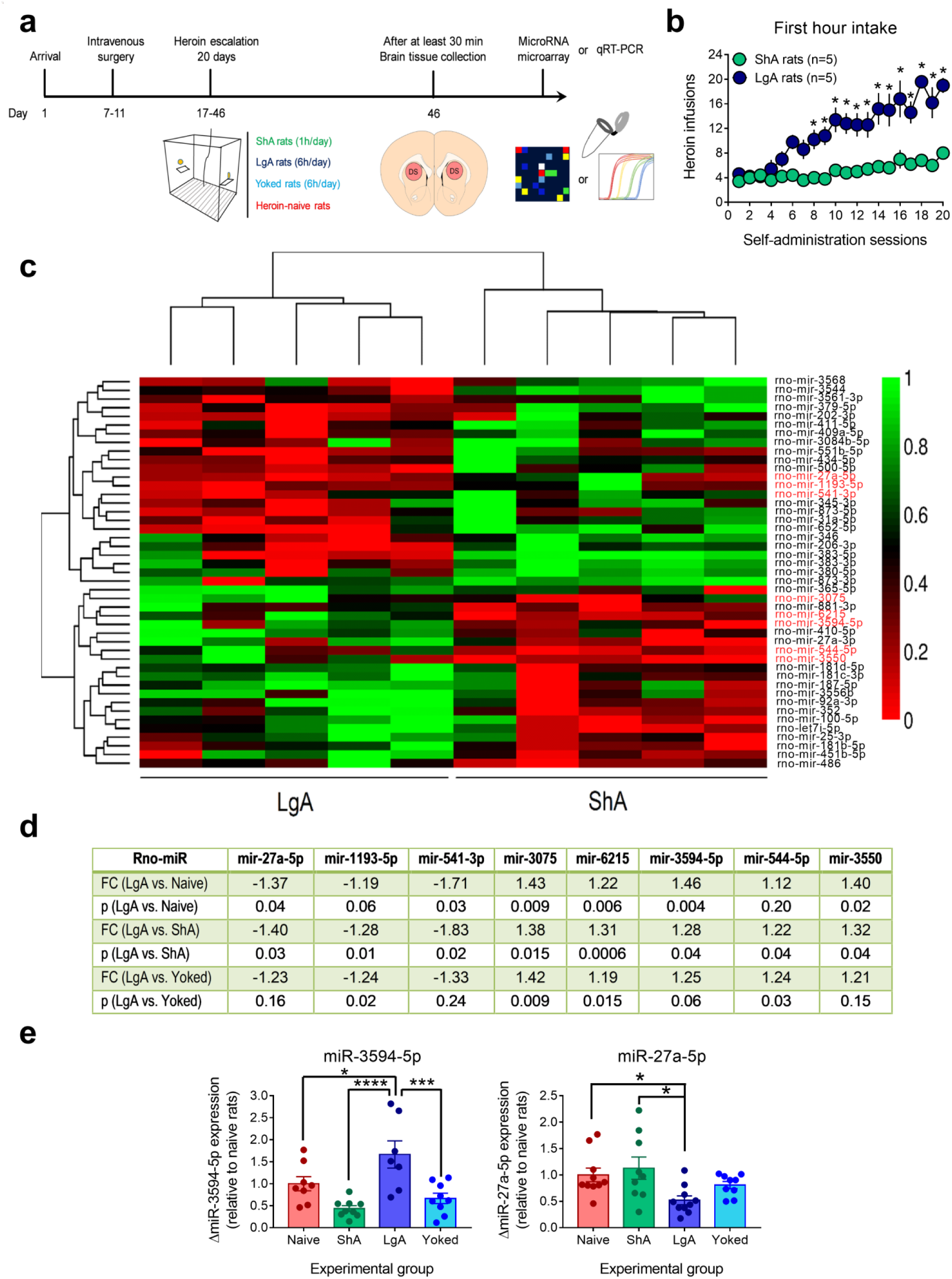
MiR-3594-5p level is increased in the dorsal striatum of extended access rats after 20 days of heroin self-administration. **a)** Timeline of the experimental procedure. **b)** Heroin infusions earned during the first hour of heroin self-administration over 20 daily sessions in restricted (ShA rats, green circles, 1h/day) and extended access rats (LgA rats, dark blue circles, 6h/day). *different from ShA rats, P<0.05, Bonferroni’s multiple-comparisons test. **c)** Heatmap and hierarchical clustering of differentially expressed miRNAs in the dorsal striatum between ShA and LgA rats after heroin self-administration (Fold change ≥ 1.2, p<0.05). Upregulated miRNAs (green) are those with increased expression levels in the striatum, whereas downregulated miRNAs are shown in red. **d)** Table summarizing the selected candidate miRNAs from the microarray data for a subsequent measure with qRT-PCR in the dorsal striatum of LgA, ShA, Yoked and Naive rats. **e)** qRT-PCR confirmed that miR-3594-5p was selectively overexpressed in the dorsal striatum of LgA rats (n=7) compared with Naive (n=8), ShA (n=9) and Yoked (n=9) rats (*,different from LgA rats, *P<0.05, ***P<0.001, **** P<0.0001, Bonferroni’s multiple-comparisons test). Striatal miR-27a-5p levels were also decreased in the LgA rats (n=10) compared with heroin-naive (n=10) and ShA rats (n=9), but not with Yoked rats (n=9) (*different from LgA rats, P<0.05; Dunn’s multiple comparison test). Data are presented as mean ± s.e.m. For qRT-PCR analyses, relative microRNA expression levels were calculated using the method of 2−ΔΔCt with snoRNA and U87 as the endogenous controls for normalization. See also Extended Data Figs. 1, 2, 3 and 4. Statistical details are included in Supplementary Table 3.

Among 8 potential miRNA candidates identified by microarray (Fig. 1c, miRNAs written in red color, Fig. 1d and Extended Data 1d, e), validation with TaqMan qRT-PCR assay confirmed the increase of miR-3594-5p in the DS of LgA rats compared with heroin-naive, ShA, LgA and LgA-Yoked rats (one way ANOVA: F_3, 29_ = 9.42, P<0.0002, Fig. 1e). We also confirmed by qRT-PCR the observed decrease of miR-27a-5p levels in the DS of LgA rats compared to ShA and heroin-naïve rats (Kruskal-Wallis ANOVA: H=10.48, P=0.0149), but not compared to LgA-Yoked rats (Fig. 1e). This result suggests that the alteration of this miRNA by extended heroin exposure is independent of operant responding. This is in accordance with the decrease of miR-27a expression reported in animals that develop antinociceptive tolerance^31^. Two specific mature miRNAs measured by microarray, miR-3075 and miR-6215 that were upregulated only in the DS in LgA rats (Extended Data Fig. 1d) were not confirmed by qRT-PCR assays and in another independent cohort of rats (Experiment 2, Extended Data Fig. 1e and Extended Data Fig. 3). In addition, no miRNA was found to be specifically altered in NAc in LgA rats (Extended Data Fig. 2b). Alterations in levels of the 4 remaining candidates, miR-1193-5p, miR-541-3p, miR-544-5p and miR-3550, were not confirmed in the DS (Extended Data Fig. 4 a-f and see Supplementary Table 3). In agreement with the microarray data, we did not observe changes in striatal miR-212 and miR-132 expression levels (Extended data Fig. 4 g-j), contrary to what were found previously with extended access to cocaine^18^.

### Striatal miR-3594-5p is a specific marker for late-stage heroin intake escalation

In order to assess whether the upregulation of miR-3594-5p in the DS is specific to extended heroin use, we trained rats to self-administer cocaine using the same differential drug access procedure (see Methods, Experiment 3, Fig. 2a and Extended Data Fig. 5a). Similarly, we analyzed microRNA profile alterations as a function of the cocaine self-administration regimen (cocaine-naive, ShA, LgA and LgA-Yoked groups, Fig. 2b and Extended Data Fig. 5b-d). We did not find any alteration in miR-3594-5p expression in the DS in LgA rats (two-way repeated measures ANOVA: Group x Session: F_19, 152_ = 4.36, P<0.0001, Fig. 2b and Extended Data Fig. 5d, e), confirming that this upregulation was specific to extended heroin use. We identified 2 miRNAs (let-7f-1-3p and miR-211-5p) and 3 miRNAs (miR-125b-5p, miR-193a-3p and miR-3588) that were specifically downregulated or upregulated, respectively, in the DS in LgA rats (see the Venn Diagram, Fig. 2c). Note that miR-125b upregulation in the DS has been previously reported after cocaine self-administration in rats^14, 18^. Against all expectations, expression levels of miR-212 and miR-132 from the same cluster, measured by microarray analysis, were not altered in the DS in LgA rats (Extended Data Fig. 5e), a negative result confirmed by qRT-PCR assay (Fig. 2d-g). Finally, no accumbal miRNA profile was specifically altered in LgA rats (Extended Data Fig.5f).

**Fig. 2.**
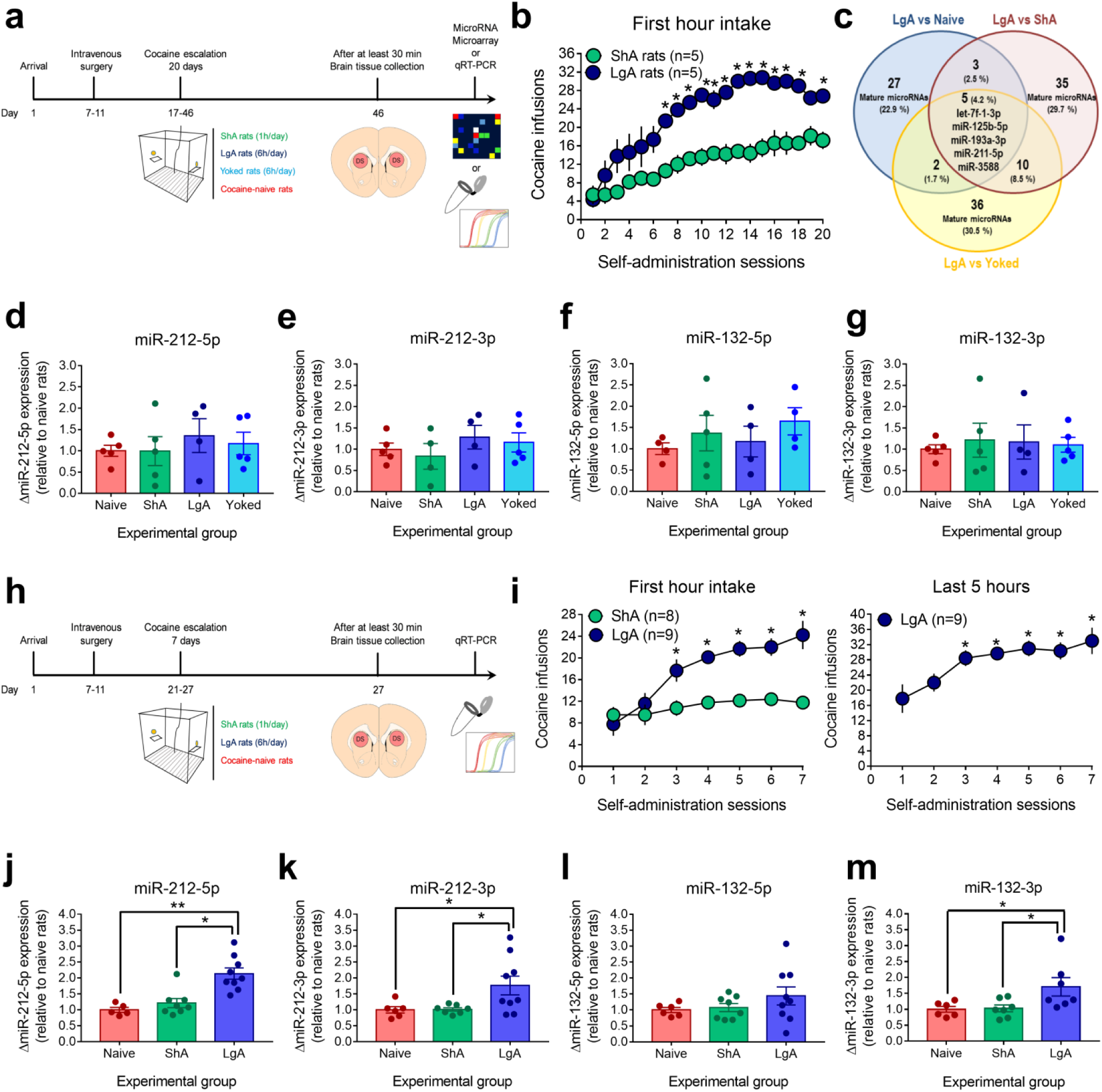
MiR-212 overexpression in the early stage but not late stage of cocaine escalation. **a)** Timeline of the experimental procedure. **b)** Cocaine infusions earned during the first hour of cocaine self-administration over 20 daily sessions in restricted (ShA rats, green circles, 1h/day) and extended access rats (LgA rats, dark blue circles, 6h/day). *different from ShA rats, P<0.05, Bonferroni’s post hoc test. **c)** Venn diagram representing the overlap of mature miRNAs significantly differentially expressed in the dorsal striatum according to the drug history (n=5 rats per group). Two miRNAs (rno-let-7f-1-3p, rno-mir-211-5p) were downregulated and 3 miRNAs (rno-mir-125b-5p, rno-mir-193a-3p, rno-mir-3588) overexpressed only in LgA rats compared with the other groups. **d-g)** Taqman qRT-PCR confirmed that striatal miR-212 and the closely miR-132 from the same cluster were not upregulated in the dorsal striatum of LgA rats with 6h daily access to cocaine compared with the other groups (one-way ANOVA, n=4-5 per group). **h)** Schematic representation of timeline for experimental design to measure miRNA levels following a short period of differential access to cocaine. **i)** Number of cocaine infusions earned by both groups during the first hour of self-administration sessions and by LgA rats during the last remaining 5 hours over the 7 daily sessions. *different from ShA rats, P<0.05; Bonferroni’s multiple-comparisons test. In LgA rats, cocaine intake also progressively increased during the 5 remaining hours (*different from the first session, *P<0.05; Bonferroni’s post hoc test). **j-m)** Taqman assay verified that striatal miR-212-5p, miR-212-3p and miR-132-3p levels were increased in extended access rats (n=9) but not in restricted (n=8) and cocaine-naïve rats (n=6) (*different from LgA rats, *P<0.05, **P<0.01, ***P<0.001, ****P<0.0001, Dunn’s or Bonferroni’s multiple comparisons test). Data are presented as mean ± s.e.m. For qRT-PCR analyses, relative microRNA expression levels were calculated using the method of 2−ΔΔCt with snoRNA and U87 as the endogenous controls for normalization. See also Extended Data Fig. 5. Statistical details are included in Supplementary Table 3.

To reconcile our observations with previous reports^18, 19^, we hypothesized that miR-212/miR-132 in the DS could be altered at early but not at late stages during cocaine intake escalation. To test this hypothesis, we measured miR-212 expression in the DS following only 7 days of differential access to cocaine self-administration (Experiment 4, Fig. 2h, i), when LgA rats just started to show escalation of drug intake (1^st^ hour: two-way repeated measures ANOVA: Group x Session: F_6, 90_ = 9.92, P<0.0001; 5 hours: one-way repeated measures ANOVA: F_2.265, 18.12_=10.71, P=0.0006, Fig. 2i). Consistent with our hypothesis, miR-212 and miR-132 expression levels were indeed upregulated, 1.5 to 2-fold in the DS in LgA rats (Kruskal-Wallis ANOVA: j, H=13.68, P<0.0001; one-way ANOVA: k, F_2,19_=4.42, P=0.0266; l, F_2,20_=1.28, P=0.2987, NS; m, F_2,17_=4.39, P=0.0291; Fig. 2j-m).

Because of this finding, we also evaluated whether the expression of miR-3594-5p in the DS is also stage-specific during extended heroin access. For that purpose, rats were given differential access to heroin self-administration until we obtained a significant difference in first-hour heroin intake between ShA and LgA rats for at least 3 daily sessions (two-way repeated measures ANOVA: Group x Session: F_11, 198_ = 1.91, P=0.04; Experiment 5, Fig. 3a,b). Note that during this early stage of heroin intake escalation, LgA rats already increased significantly their heroin intake during the last 5 hours compared with the first session (one-way repeated measures ANOVA: F_2.253, 18.03_=3.55, P=0.046, Fig. 3c). Following 12 days of differential access to heroin self-administration, there was no alteration in miR-3594-5p (Fig. 3d) and miR-27a-5p (Fig. 3e) expression levels in the DS. Similarly, there was also no alteration in miR-212 (Fig. 3f, g) and miR-132 (Fig. 3h,i) (see Supplementary Table 3 for statistical details). Taken together with findings from the first experiment, these results suggest that heroin-specific alterations in striatal miRNA emerge only late during extended heroin access after escalation has plateaued.

**Fig. 3.**
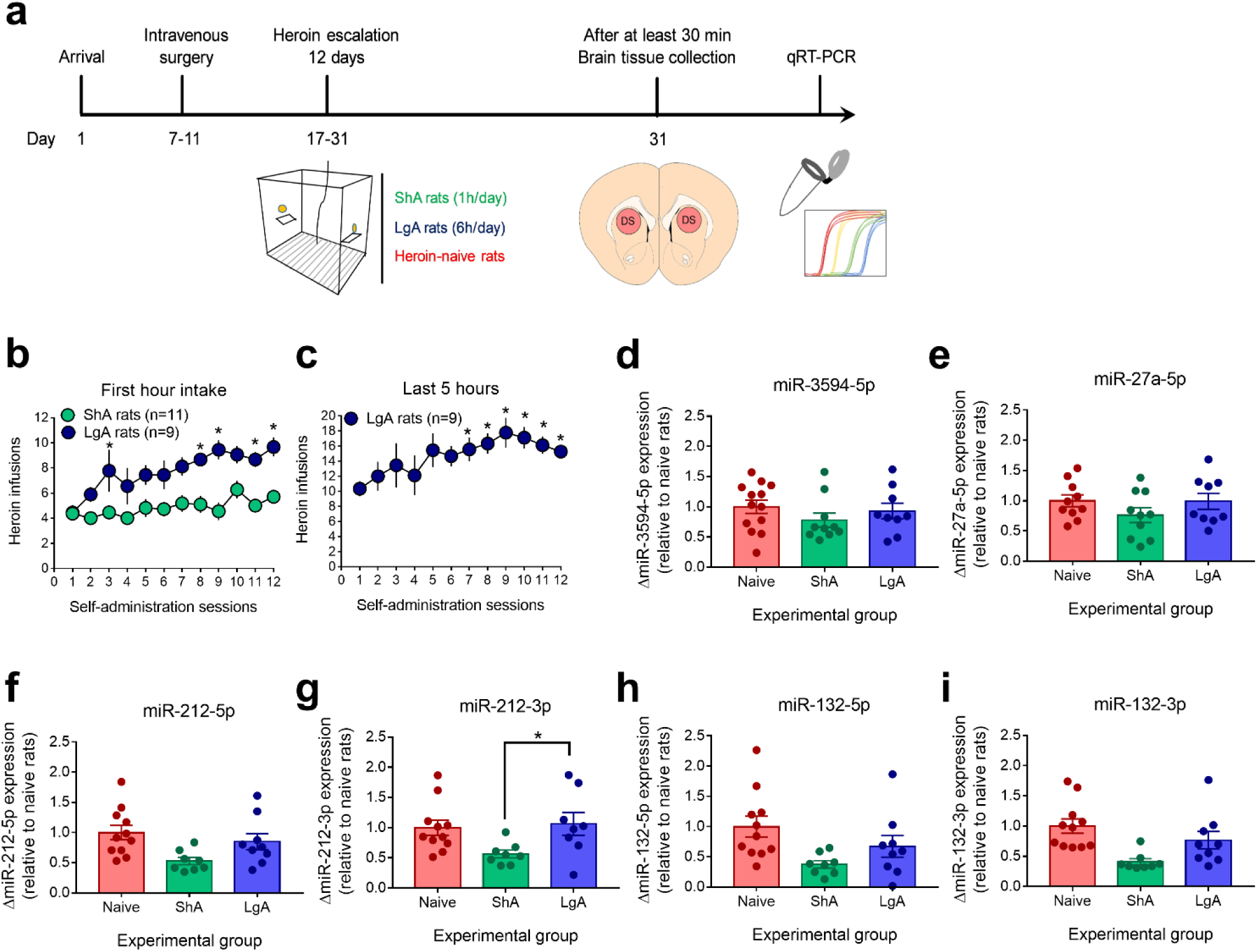
miR-3594-5p is not upregulated in the dorsal striatum of extended access rats during the early stage of heroin escalation. **a)** Timeline of the experimental procedure. **b)** Effects of access time to heroin on the number of heroin injections during the first hour of each self-administration session over 12 daily sessions (*different from ShA rats, P<0.05, Bonferroni’s post hoc test). **c)** Effects of access time to heroin on the number of heroin injections during the last 5 hours of each self-administration session over 12 days (*different from the first session, *P<0.05, Bonferroni’s post hoc test). **d-e)** During the early phase of heroin escalation, we observed no change in miR-3594-5p and miR-27a-5p levels in the dorsal striatum of LgA rats (n=9) compared with ShA (n=11) and heroin-naïve rats (n=13). **f-i)** Taqman qRT-PCR confirmed that striatal miR-212 and the closely miR-132 from the same cluster were not upregulated in the dorsal striatum of LgA rats compared to both heroin-naïve and ShA groups (one way ANOVA: (f) F_(2, 25)_ =4.32, P=0.0244; (g) F_(2, 24)_ = 3.72, P=0.0393; (h) F_(2, 25)_ = 3.98, P=0.0315; Bonferroni’s multiple-comparisons test, *different from LgA rats, P<0.05; and Kruskal-Wallis ANOVA: (i), H=12.08, P=0.0024; Dunn’s multiple-comparisons test). Data are presented as mean ± s.e.m. For qRT-PCR analyses, relative microRNA expression levels were calculated using the method of 2−ΔΔCt with snoRNA and U87 as the endogenous controls for normalization. Statistical details are included in Supplementary Table 3.

### Top canonical pathways enriched by predicted targets of striatal miR-3594-5p

To identify gene targets for miR-3594-5p, we imported our miRNA microarray data into the Ingenuity Pathway Analysis Software to analyze the list of predicted targets and related functional pathways. Overall, we found 522 predicted targets for miR-3594-5p (Supplementary Table 4). In addition, miR-3594-5p was predicted to regulate genes associated with 19 significantly enriched canonical pathways (Extended Data Fig. 6a) including axonal guidance signaling (-log(p-value)=1.75), Toll-like receptor signaling (-log(p-value)=1.42) and dopamine-DARPP32 Feedback in cAMP signaling (-log(p-value)=1.31). In another independent cohort of rats (Experiment 6, Extended Data Fig. 7a), we decided to test the validity of 12 potential target transcripts of interest associated to the axonal guidance signaling pathway using qRT-PCR assays (Extended Data Fig. 6b). However, we did not observe any decrease in the expression of these transcripts in the DS in LgA rats (see Supplementary Table 3 for statistical details) (First hour intake: two-way repeated measures ANOVA: Group x Session: F_19, 342_ = 4.83, P<0.0001; 5 last hours for LgA rats: on way ANOVA: F_3.775_,_33.97_=12.33, P<0.0001; Active lever presses: Session x Group Interaction: F_19,342_=2.54, P=0.0005; Inactive lever presses: Session x Group Interaction: F_19,342_=1.004, P=0.4556; Fig. 4d and Extended Data Fig. 7b-e). Amongst the 522 predicted target transcripts, we determined that 38 targets were predicted by at least 2 of the 3 databases used (TargetScan, MiRDB and Diana-microT-CDS) and only 11 targets were predicted by all the databases: *Bbs4, Cdip1, Cdipt, Chac1, Chchd4, Csf1, Grb10, Nras, Rpl41, Sez6* and *Usp42* (see Supplementary Table 4). Importantly, these 11 transcripts were predicted to be specifically targeted by miR-3594-5p, but not by miR-27a-5p. Of the 11 target genes, *Sez6* was the transcript with the highest prediction scores (followed by *Grb10* and *Nras*) in two of the three algorithm databases (Fig. 4a, Table), giving us more confidence in this prediction.

**Fig. 4.**
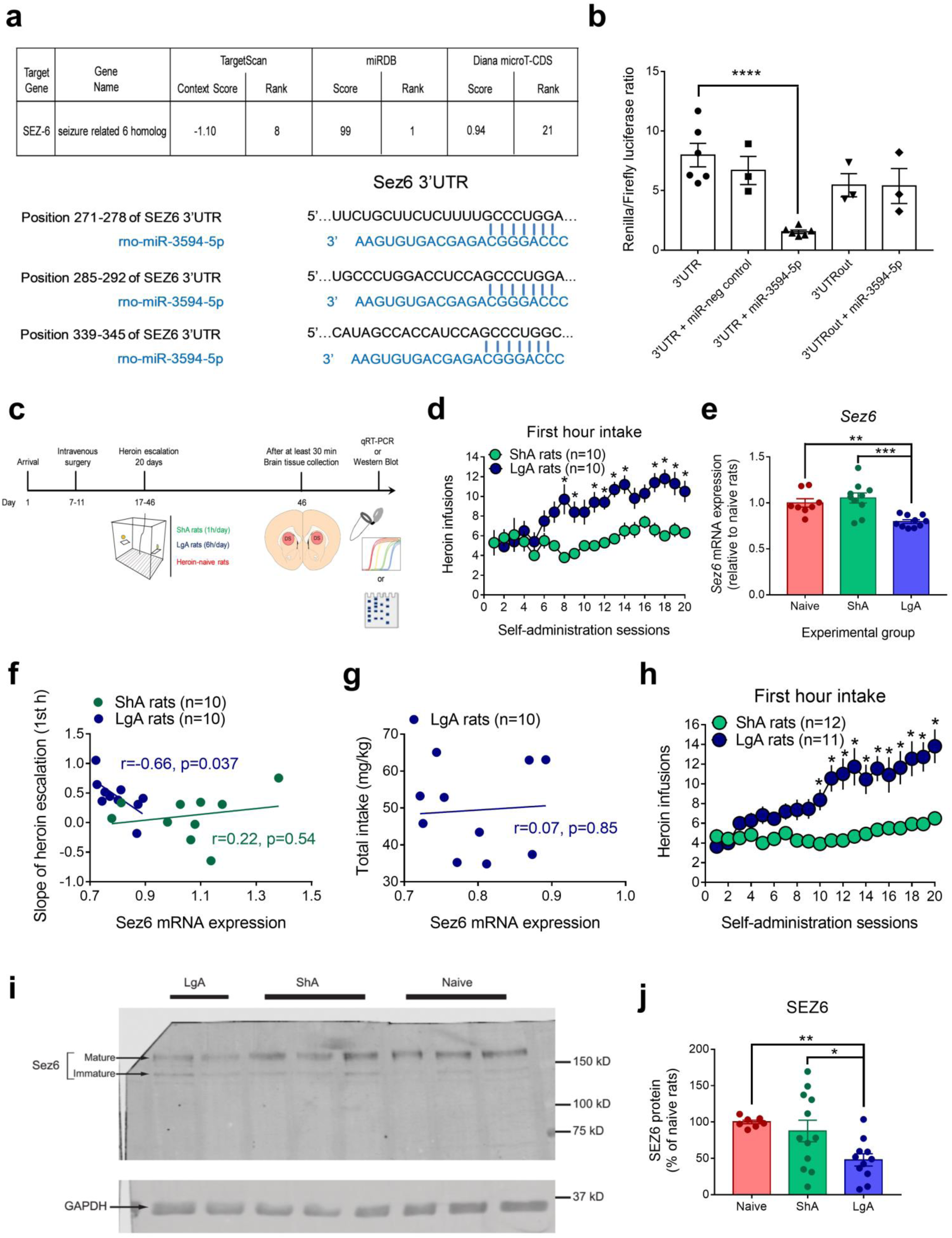
miR-3594-5p inhibits mRNA and protein expression of SEZ-6 in the dorsal striatum of rats with escalating patterns of heroin intake after 20 days of extended access to the drug. **a)** In silico prediction and binding of miR-3594-5p to *Sez6*. Prediction scores for *Sez6* predicted by 3 databases are included in the table. The binding pairs are indicated by the blue bars between the miRNA and the mRNA. **b)** In vitro dual luciferase reporter assays of miR-3594-5p-*Sez6* mRNA interaction. HEK239 cells were co-transfected with *Sez6* 3’UTR, *Sez6* 3’UTR and miRNA negative control mimic, *Sez6* 3’UTR and miR-3594-5p, 3′-UTRout *Sez6* construct, 3’UTRout and miR-3594-5p in dual-luciferase assays (*different from 3’UTR, ****P<0.0001, Bonferroni’s multiple comparisons test). See Material and Methods part for more details about luciferase assay constructs. **c)** Timeline of the experimental procedure for *Sez6* mRNA (Experiment 6) and SEZ6 protein (Experiment 7) quantifications. **d)** Extended access to heroin produced escalation of intake over time (Experiment 6). *different from ShA rats, P<0.05, Bonferroni’s multiple-comparisons test. **e)** Sez6 mRNA expression is downregulated in the dorsal striatum of LgA rats (n=10) compared with heroin-naive (n=8) and ShA rats (n=10) (*different from LgA rats, *P<0.05, ***P<0.001, Bonferroni’s multiple comparisons test). Relative *Sez6* mRNA expression levels were calculated using the method of 2−ΔΔCt with *Rplpo* and *Hmbs* as the endogenous controls for normalization. See also Extended Data Fig.7 for more details about *Sez6* mRNA quantifications. **f)** Pearson correlations between the slope of escalation in heroin intake for each individual during the first hour of self-administration access and the *Sez6* mRNA expression (relative to heroin-naive rats) in ShA (green, n=10) and LgA (blue, n= 10) rats (see Material for more details). **g)** Pearson correlation between the cumulative heroin intake (mg/kg of body weight) exhibited by LgA rats and the *Sez6* mRNA expression (relative to heroin-naive rats). **h)** Effect of access time to heroin on the number of heroin injection during the first hour of each self-administration session before protein quantification (Experiment 7). **i)** Representative image of the full-length western blot gels and **j)** mature SEZ6 protein expression normalized to GAPDH following differential access to heroin for 20 days. LgA rats (n=11) showed a significant decrease in SEZ6 level in the dorsal striatum compared with heroin-naive (n=8) and ShA animals (n=12) (*different from LgA rats, *P<0.05, **P<0.01; Dunn’s multiple comparison test). See also Extended Data Fig. 8 for more details about protein quantifications. Data are presented as mean ± s.e.m. Statistical details are included in Supplementary Table 3.

### Striatal SEZ-6 is selectively inhibited by miR-3594-5p in LgA rats

As already reported, *Nras* expression levels were unchanged in the DS in LgA rats compared to the other groups (one-way ANOVA: F_2,27_=0.4938, P=0.6157, Extended Fig. 6b). Similarly, no change was observed in *Grb10* expression in the DS in LgA rats compared to heroin-naive rats (one-way ANOVA: F_2,26_=6.575, P=0.0049, Naïve vs. LgA: P>0.9999, ShA vs. LgA: P=0.0085, Bonferroni’s post hoc test, Extended Data Fig. 7f). Most notable was *Sez6* (Seizure related 6 homolog), a transcript which is translated into an N-glycosylated type I transmembrane protein, expressed in neuronal cells and abundant in the striatum of adult rodents^32^. SEZ6 was of particular interest because it has been demonstrated to play a critical role in proper dendritic branching, synapse formation and transmission, and LTP^33, 34^. Further, *in silico* analysis predicted that miR-3594-5p would exert its action by binding to the 3’UTR of *Sez6* mRNA at 3 potential sites to facilitate its degradation and/-or inhibit its translation (Fig. 4a). To probe the direct interaction between miR-3594-5p and the 3’UTR sequences of *Sez6* mRNA, we employed a dual luciferase reporter assay (Fig. 4b, see Methods for plasmid constructs). Co-transfection of the plasmid containing the wild-type *Sez6* 3’UTR with miR-3594-5p into HEK293 cells strongly reduced the luciferase activity (one way-ANOVA: F_4,16_=9.425, P=0.0004; 3’UTR vs. 3’UTR + miR-3594-5p: P<0.0001, Bonferroni’s post hoc test). This reduced activity was specific to miR-3594-5p because co-transfection with the microRNA negative control did not affect the luciferase activity (3’UTR vs. 3’UTR + miR-neg control: P>0.9999, NS). Moreover, luciferase activity was not affected when cells were co-transfected with the plasmid containing the 3’UTRout and the miR-3594-5p mimic (3’UTR vs. 3’UTR + miR-3594-5p: P=0.2757, NS). Note that 3’UTRout did not, by itself, affect the luciferase activity (3’UTR vs. 3’UTRout: P=0.3029). Overall, these results confirmed experimentally that miR-3594-5p directly interacts with the 3’UTR of *Sez6* mRNA.

Next, we aimed to verify whether *Sez6* mRNA expression was down-regulated in LgA rats following 20 days of heroin self-administration (Fig. 4d). As expected, *Sez6* mRNA level was significantly decreased by 20 % in the DS of LgA rats compared with heroin-naïve rats and by 28% when compared with ShA rats (one-way ANOVA: F_2,26_=9.85, P=0.0007, LgA vs. Naïve: P=0.0126, LgA vs ShA: P=0.0004, Bonferroni’s post hoc test Fig. 4e). Importantly, the individual slope of escalation in heroin intake during the first hour of self-administration session in LgA rats – a measure of escalation severity – was negatively correlated (r=-0.66, P=0.037, Pearson’s correlation) with striatal *Sez6* mRNA expression levels, while no significant correlation was found in ShA rats (r=0.22, P=0.55) (Fig. 4f). Thus, the larger the decrease in *Sez6* mRNA expression levels in the DS in LgA rats, the higher the severity of heroin intake escalation. Interestingly, there was no correlation between *Sez6* expression levels in the DS in LgA rats and total amount of heroin consumed during the experiment (r=0.07, P=0.85, Fig. 4g). Although not significant, LgA rats with more severe escalation also tended to have a higher level of miR-3594-5p in the DS (r=0.63, p=0.089, NS, Pearson’s correlation). In the same individuals and as expected, the striatal *Sez6* mRNA expression level was highly negatively correlated with the miR-3594-5p expression level (r=-0.78, p=0.0396, Pearson’s correlation). Finally, we measured SEZ6 protein expression in the DS in another independent cohort of rats (Experiment 7, Fig. 4c) submitted to 20 days of differential access to heroin (First hour intake: two-way repeated measures ANOVA: Group x Session: F_19, 399_ = 7.72, P<0.0001; last 5 hours for LgA rats: one-way ANOVA: F_2.704_,_27.04_=9.75, P=0.0002; Active lever presses: Session x Group Interaction: F_19,399_=3.91, P<0.0001; Inactive lever presses: Session x Group Interaction: F_19,399_=1.15, P=0.3002; Fig. 4h and Extended Data Fig. 8a-d). Polyclonal anti-SEZ6 antibody detected two bands: a main band at around 170 kDa and another weaker band at around 150 kDa (Fig. 4i for representative image of the full-length western blot gels). As previously reported, these bands are specific to the SEZ6 polyclonal antibody^33, 35^. It has been already demonstrated that the 170 kDa band corresponds to SEZ6 containing complex glycosylated sugars (i.e., mature SEZ6), whereas the 150 kDa band is a SEZ6 form containing only high mannose sugars (i.e., immature SEZ6)^35, 36^. We found that mature SEZ6 protein expression was reduced by approximately 52% relative to the heroin-naïve rats and by 40% relative to ShA rats (Kruskal-Wallis ANOVA: H=8.969, P=0.0113, Naïve vs. LgA: p = 0.0044, ShA vs. LgA: P=0.0309, Dunn’s post hoc test, Fig. 4j), in accordance with the data obtained on *Sez6* mRNA.

## DISCUSSION

Our study reveals that miR-3594-5p level was upregulated in the DS following extended but not restricted heroin access, a modification not seen following extended cocaine access. This upregulation appeared at late, but not early stages of heroin intake escalation when excessive consumption plateaued. This upregulation was not observed in yoked controls, indicating a specific association with excessive self-administration of heroin. miR-3594-5p bound directly to the 3’UTR region of *Sez6* transcript and inhibited its expression, resulting in the decrease in the mature form of the translated SEZ6 protein in the DS of LgA rats. A negative correlation was observed between *Sez6* expression and severity of heroin intake escalation. Inversely and as expected, miR-3594-5p levels tended also to be positively correlated with escalation severity. Overall, these findings suggest that inhibition of striatal SEZ6 protein by miR-3594-5p is a molecular marker for late-stage heroin intake escalation.

### Intake escalation-related miRNA regulations are drug-specific

Expression of some miRNAs, with target genes related to synaptic plasticity, are altered by multiple classes of drugs of abuse, such as the miR-212/132 cluster, let-7, miR-9 and miR-124^37^. However, some other miRNA alterations seem to be specific to a single substance of abuse or a pharmacological class of drug. In the present study, we confirmed that striatal changes in miRNAs levels involved in the escalation process were specific to the type of drug available for self-administration (cocaine *versus* heroin). Extended access cocaine self-administration resulted in an upregulation of miR-212/miR-132 expression in the DS at early stages of cocaine intake escalation, consistent with previous research^18^. In contrast, no change in expression of these miRNAs was observed with heroin, regardless of the stage of drug intake escalation. We also found an increase of miR-3594-5p expression in the DS after extended access to heroin, but not cocaine, for self-administration. This is in accordance with the fact that heroin and cocaine trigger different long-lasting neurobiological adaptations within cortico-striatal circuits^38^ and that the pharmacological and behavioral factors underlying escalated drug overconsumption largely vary between these two drugs^39^. However, because of their pleiotropic nature, each miRNA could potentially target hundreds or thousands of mRNAs^40^. We thus cannot exclude that different miRNAs are altered during escalation of heroin and cocaine consumption but that they regulate the same target transcripts. Nevertheless, we found that miR-125b-5p, miR-193a-3p and miR-3588 were upregulated and let-7f-1-3p and miR-211-5p downregulated in the DS of rats, which had previously escalated their cocaine intake over 20 days. None of these miRNAs has *Sez6* as a predicted target. Hence, it seems that the involvement of miR-3594-5p/*Sez6* axis in the escalation process is specific to heroin.

### Spatiotemporal specificity of miRNA alterations in escalation of cocaine and heroin intake

Heroin and cocaine induced highly specific spatial and temporal changes in the expression pattern of miRNAs. Consistent with previous reports^18, 19^, we found that miR-212/miR-132 were upregulated in the DS, but not in the NAc, of LgA rats only during early stages of cocaine intake escalation. However, we found for the first time that this alteration vanished when escalation of cocaine intake reached its plateau and was maintained. We hypothesized that, once excessive cocaine intake has developed, the protective function of striatal miR-212 would no longer be needed or would be relayed by another molecular mechanism. Interestingly, it was previously reported that 14 days of short access to cocaine (2h per day) increased both miR-212 and miR-132 levels in the DS of rats and this effect endured at least 10 days after rats had been withdrawn from the drug during extinction training^41^. In our conditions, we did not find any increase of miR-212/miR-132 in ShA rats following 20 days of cocaine self-administration (1h per day). More importantly, the fact that we no longer observed changes in levels of these miRNAs in the DS of LgA rats following 20 days of differential access to cocaine suggests that other molecular neuroadaptations were engaged resulting directly or indirectly from excessive cocaine consumption. Another possible interpretation, suggested by previous studies^42, 43^, is that miR-212 and miR-132 are involved at all the different stages of the escalation process but differently according to the subregions of the DS. miR-212/132 levels would be increased within the dorsomedial striatum (DMS) early in the escalation process, whereas their expression levels would be both decreased and increased within the DMS and the dorsolateral striatum (DLS), respectively at the late stage. These temporal dynamics of miR-212/132 expression would reflect the early engagement of DMS and the gradual devolution of control from the DMS to DLS during the transition from drug use to addiction. As we measured miRNA alterations in the whole dorsal striatum, the apparent lack of change at late stages of cocaine intake escalation could result from opposite miRNA alterations within the two subregions. Further research is required to fully clarify the precise temporal pattern of miR-212/132 expression in the DMS and DLS during the transition to cocaine addiction. Whatever the hypothesis considered, however, miR-212/132 alterations critically involved in cocaine escalation are spatiotemporally-regulated. Contrary to the upregulation of miR-212/miR-132 measured from the very beginning of the development of cocaine escalation, the upregulation of striatal miR-3594-5p was observed at late stages of heroin intake escalation in the DS, but not the NAc, when excessive consumption plateaued. This finding suggests that this change could be a result of the escalation process and may be involved in the maintenance of heroin excessive consumption. This is of particular interest in the context of translational biomarker research for medication development, given that people with substance abuse disorders often consult once excessive drug use has already been established.

### Potential role of Sez6 in escalation of heroin consumption

Because miR-3594-5p is a translational repressor which acts by binding to a specific sequence in the 3’UTR of its target mRNAs, we sought to identify target genes of miR-3594-5p that may be involved in excessive heroin-taking behavior. We found that SEZ6 expression is selectively inhibited by miR-3594-5p within the DS only in heroin extended-access rats. Interestingly we observed that LgA rats with higher severity of heroin intake escalation (or greater individual escalation slope) exhibited a higher decrease in *Sez6* mRNA expression levels. SEZ6^44, 45^ is a little-studied N-glycosylated type I transmembrane protein, widely expressed at the somatodendritic surface of neurons and abundant in the striatum of adult rodents^32, 46^. SEZ6 has been shown to have fundamental functions in the nervous system, including in dendritic branching, synapse formation and transmission (particularly excitatory synaptic connectivity), and LTP^33, 34^. Recently, SEZ6 was reported to bind the kainate receptor subunits GluK2/3 (GRIK2/3), which are highly expressed in the DS^47^, and promotes their intracellular trafficking, glycosylation, surface localization and ion channel activity in CA1 pyramidal neurons^48^. Kainate receptors participate in short-term plasticity, in LTP and LTD processes^49^. In addition, GRIK2 and GRIK3 are linked to obsessive compulsive disorder in humans^50^. Knockout mice, in which all five kainate receptor subunits are ablated, display compulsive and perseverative behaviors associated with corticostriatal dysfunction^51^. Overall, these results argue that excessive heroin consumption induces an increase of miR-3594-5p level in the DS, which in turn decreases SEZ6 expression. SEZ6 inhibition would alter kainate receptor signaling (presumably by decreasing kainate receptors on the neuron surface and reducing kainate-evoked currents), which may play a role in the maintenance of excessive, uncontrolled and perseverative heroin taking via its action on long-term synaptic plasticity. The present findings do not allow us to determine whether the miR-3594-5p-mediated downregulation of *Sez6* mRNA results from heroin escalation and contributes to maintain excessive use. Further research is needed to validate the causal role of miR-3594-5p/*Sez6* interaction in the maintenance of escalated heroin use.

### Conclusion and Potential implications

It is important to note that miR-3594-5p is specific to rat and is not conserved in humans. However, the *Sez6* gene is highly conserved in humans, suggesting that it may be implicated also in excessive heroin use in humans. In conclusion, we have characterized a new miR-3594-5p/*Sez6* axis within the DS associated with escalating heroin use in rats, discovered a new potential gene target conserved in humans and suggested future directions for research on the mechanisms involved in escalation of heroin use.

## METHODS

### Subjects

237 young adult Wistar male rats, weighing 200-225 g at their arrival in the laboratory, were purchased from Janvier Labs (France) and randomly housed in groups of two in home cages placed on a ventilated cage rack (Tecniplast, France). We maintained the rats in a light- (12-h reverse light-dark cycle, light off at 8 am) and temperature-controlled vivarium (22°C). We provided a transparent rectangular plastic red toy and 4 squares of cotton inside each cage to promote animal well-being. Standard laboratory chow and tap water were freely available in the home cages during the whole duration of behavioral procedures. All behavioral testing occurred during the dark phase of the light-dark cycle. All experiments were carried out in accordance with the European (2010/63/UE, 20 October 2010) and French (decree n ° 2013-118, 2013) directives, and approved by the local ethics committee (Comité d’Ethique en matière d’Expérimentation Animale de Paris Descartes, CEEA 34). We excluded 49 rats from the experiments due to loss of catheter patency (n=26), illness (n=9) and failure to acquire the self-administration behavior (n=14).

### Intravenous surgery

We transiently anesthetized rats with isoflurane gas (induction: 4%) to inject a mixture of ketamine and xylazine (80 mg/kg, Virbac and 10 mg/kg, Bayer; i.p.). Anesthetized rats were implanted with indwelling silastic catheter into the right jugular vein, as previously described^52^. The distal end of the catheter exited the skin in the middle of the back via a metal guide cannula. After surgery, rats were allowed to recover for a period of at least 7 days before starting behavioral testing. After surgery and during the whole duration of the experimental procedures, catheters were flushed daily with a prophylactic treatment of an antibiotic (Amoxicillin, 90 mg/kg/day) dissolved in a sterile heparinized saline NaCl 0,09% (B. Braun – héparine choay® 240 UI/ml). If necessary, catheter patency was checked by administering 0.15 ml of the short-acting non-barbiturate anesthetic Etomidate (Hypnomidate 2 mg/ml, Janssen) through the catheter.

### Self-administration chambers

Rats were trained in 12 identical self-administration chambers (40 x 30 x 36 cm, Imetronic, Pessac, France) located in a dimly lit room by red lights and enclosed in a ventilated and sound-attenuated wooden enclosure with a white noise generator (50 dB) to minimize distracting visual pollution and noise. Each chamber was constituted of two operant panels facing each other, in the middle of which was mounted a retractable and motorized lever located 7 cm above the grid floor. A light diode was mounted above each lever (white light diode of 1.2 cm of outer diameter). During self-administration sessions, presses on the left active lever controlled the activation of the syringe pump (Imetronic, France) placed outside, on the top of the enclosure. This pump delivered a drug infusion in a volume of 37 µl over a period of 1 s via a Tygon tubing (Cole Parmer Instrument) protected by a stainless steel spring. The cannula connector (Plastic One, VA, USA) on the back of the animal was connected to a single-channel liquid swivel via the Tygon tubing. We then connected the liquid swivel to a 20-ml syringe through tubing. Presses on the right inactive lever were not reinforced by the drug. Two sound generators were located on the top of the cage. Experiment piloting, data collection and data storage were controlled by a computer via an interface and specific software (Imetronic, France).

### Differential access to intravenous heroin self-administration

In Experiment 1, we used the well-established extended-access model^1, 28^ to determine the impact of access time to heroin for self-administration on miRNA profiling in the NAc and DS by using miRNA 4.0 Affymetrix GeneChips. One week after surgery, rats were first habituated to the conditioning chambers for one session of 3 hours before starting the self-administration training. During this session, both levers were retracted and the white noise (50 dB) was turned on. The following day, all rats were tested for 1h-session of intravenous heroin self-administration (15 µg per unit dose delivered in a volume of 74 µl over 2 seconds) under a fixed-ratio 1 with 20-s timeout reinforcement schedule. The delivery of heroin was signaled by the illumination of the cue light above the lever for 20s and by the delivery of a 5s-tone (3000 Hz, 60 dB). This screening phase was a reliable method to balance the groups on the basis of mean body weight and mean number of heroin injections, as shown previously^29, 53^. During the subsequent escalation phase, one group had access to heroin for 1h/day (Short Access rats, ShA rats, n=5) and another group for 6h/day (Long Access rats, LgA rats, n=5). During the 5 remaining hours, the unit dose available was increased to 60 µg/injection to promote heroin intake escalation (by increasing the injection volume to 296 µl over 8 seconds), as previously described^29, 54^. Active lever presses led to the delivery of heroin and the compound tone-light cue, while inactive lever presses had no programmed consequence. We trained rats during 20 daily self-administration sessions (5 days/week) and tested two additional control groups: heroin-naïve (n=5) and LgA-Yoked (n=5) groups. Heroin-naïve rats underwent all experimental procedures (surgery, antibiotic treatment, housing…), except that they were never exposed to heroin or any operant contingency. We randomly assigned rats to the heroin-naïve group at the beginning of the experiment. Finally, each rat from the Yoked group was matched with an LgA rat and received non-contingently (i.e., independently from their behavior) the heroin doses that the latter obtained by self-administration (with the associated tone-light cues).

In Experiment 2, we trained rats to self-administer heroin in the same conditions as those described in Experiment 1 to confirm more quantitatively some of the relevant changes observed in miRNA levels by using TaqMan microRNA assay-based qRT-PCR (heroin-naïve, ShA, LgA and Yoked rats, n=10 rats /group). Indeed, qRT-PCR is considered as the gold standard for gene expression quantification^55^.

In Experiment 5, we trained 37 rats to self-administer heroin using the same extended access escalation model but for fewer days (ShA and LgA rats: n=12; Naive rats: n=13). To evaluate miRNA profiling within the DS at the early stage of heroin intake escalation, we trained rats for 12 days before stopping the drug self-administration procedure. In this condition, heroin consumption of LgA rats during the first hour was significantly higher than those of ShA rats for at least 3 days.

In Experiments 6 and 7 (target mRNAs and SEZ6 protein quantification studies, respectively), the protocol for heroin intake escalation used in Experiment 1 was applied, except we tested only the 3 main experimental groups (Naive: n=11 and n=12, ShA: n=11 and n=12, LgA: n=12 and n=12, for Experiments 6 and 7, respectively).

### Differential access to intravenous cocaine self-administration

All experimental conditions were kept identical to those described for heroin self-administration training, except that the unit doses of cocaine were 250 µg/infusion for the first hour (delivered in a volume of 74 µl over 2 seconds) and 750 µg/infusion for the 5 remaining hours (delivered in a volume of 222 µl over 6 seconds), as previously described^56^. We trained rats for 20 sessions (5 days/week, Experiment 3) or 7 consecutive sessions (Experiment 4).

In Experiment 3, the animals were given differential access to cocaine to determine the impact of access time to cocaine on miRNA profiling in our experimental conditions (Cocaine-naïve, ShA, LgA and LgA-Yoked groups, n=5 rats/group).

In Experiment 4, 35 rats were allowed to self-administer cocaine for only 7 consecutive days to subsequently quantify miR-212 and miR-132 levels in the DS at the early stage of cocaine escalation by using TaqMan qRT-PCR assays (ShA and LgA groups: n=12, Naive group: n=11). Data from the literature had already shown that these two miRNAs were regulated in the DS following escalation of cocaine intake^18^. This experiment was an internal control in our study.

### Total RNA extraction

Rats were euthanized by decapitation following a pentobarbital injection (Doléthal®, 70-80 mg/kg, i.v., Vetoquinol) at least 30 min after the end of the last self-administration session to prevent them being under cocaine or heroin intoxication or experiencing acute withdrawal. Brains were rapidly removed, frozen immediately in chilled isopentane and stored at -80°C until dissection. A 1.3-mm-thick coronal section (bregma: 1.7-0.2 mm according to the stereotaxic coordinates^57^) containing the NAc and the DS was obtained using a brain matrix (Braintree Scientific, Inc) and placed on a dry-ice cold glass slide under RNAse free conditions. Bilateral 1.2 mm^2^ punches from the NAc and bilateral 2 mm^2^ punches from the DS were dissected using biopsy punches, collected into Eppendorf tubes, kept in dry ice and stored at -80°C until RNA extraction. Total miRNA and mRNA were extracted from tissue samples using miRNEasy Micro Kit (Qiagen, France) according to the manufacturer’s recommendations. Frozen samples were disrupted and homogenized into 700 µl of Qiazol (mixture of phenol-chloroform and guanidium thiocyanate, Qiagen) by using a rotor-stator homogenizer (TissueRutpor II, Qiagen). We purified RNA with RNeasy Micro columns (Qiagen) and performed an optional on-column DNase digestion. We confirmed the RNA purity by using a NanoDrop 1000^TM^ or Nanodrop one™ spectrophotometer (Thermo Scientific ; 1.9≤260/280≤2.1; 1.6≤A260/230≤2.2) and the RNA integrity with the Agilent 2100 Bioanalyzer (Agilent Technologies, Santa Clara, CA) combined to the BioAgilent’s RNA Integrity Number (RIN) software (RIN>8, Genom’IC platform, Institut Cochin, Paris)^58^.

### Microarray hybridization and analysis

Microarray hybridization was performed at the Genom’IC platform (Institut Cochin, Paris) using miRNA 4.0 Affymetrix GeneChips (Affymetrix, Santa Clara, CA) according to the manufacturer’s protocol. These arrays are designed to interrogate all mature miRNA sequences in miRBase Release 21, which includes oligonucleotide probe sets for 740 rat mature miRNAs and probes for 491 rat pre-miRNAs. Briefly, 260 ng of total RNA was labeled using the FlashTag Biotin RNA Labeling Kit (Affymetrix, Santa Clara, CA). Labeled samples were hybridized to the arrays at 48°C for 20h and then washed and stained with a Streptavidin-PE solution. Arrays were scanned using the GCS3000 7G to obtain raw data (cel files) and metrics for Quality Controls. Data were normalized using RMA algorithm in Bioconductor with the custom CDF vs 21 of Brain array. Statistical analysis was performed using Partek® GS V3.0 (Partek Inc., St. Louis, MO). In our experiments, no apparent outliers were detected. MiRNAs with nominal p ≤ 0.05 between two groups of rats (ShA *versus* LgA; Naive *versus* LgA; and Yoked *versus* LgA) were considered differentially expressed to LgA rats. MiRNA expression profiling datasets are available in the Gene Expression Omnibus (GEO) database (www.ncbi.nlm.nih.gov/geo/; cocaine: accession number , heroin: accession number ).

### Computational Analysis of miRNA Targets and Pathways

To predict miRNA targets and biological pathways impacted by the differentially expressed miRNAs, we combined a bioinformatics analysis using Ingenuity Pathway Analysis (IPA, Ingenuity® Systems, Redwood City, CA, USA; www.ingenuity.com) software of our microarray data with the use of 3 different prediction algorithms (TargetScan^59^ 7.2, MiRDB^60^ and Diana-microT-CDS^61^ v5.0). Molecules predicted by at least two of the three algorithms were identified as predicted targets. The IPA software predicted the targets mRNA based on different algorithms TargetScan, miRecords and Ingenuity Knowledge Base, as well as a database devoted to the indexing of experimentally supported miRNA-gene interactions (TarBase). IPA software allowed us to identify the significantly enriched canonical biological pathways potentially influenced by target genes of differentially expressed miRNAs.

### cDNA synthesis and quantitative TaqMan RT-PCR

For TaqMan analysis of miRNA expression, we reverse transcribed RNA into cDNA in an RNAse free environment using the RT TaqMan® MicroRNA Reverse Transcription (RT) kit (Applied Biosystems, ABI) with the miRNA-specific commercial primers (ABI, Thermofisher, see Supplementary Table 1 for the list of primers used). We used small nuclear RNA (snoRNA) and U87 as endogenous controls for miRNA, validated among 4 candidates (U7, 4.5SRNA, U87 and snoRNA) in our experimental conditions using GeNorm algorithm in qBasePlus^62^. Extracted total RNA was diluted to an initial concentration of 2 or 3ng / μl. The protocol followed the manufacturer’s recommendations with the exception of using 2 μl for each RT primer (ABI) in a 10 μl total reaction volume (i.e., 3.3 μL of extract RNA, 0.1 μL of 100 mM dNTPs, 0.67 μL of MultiScribe^TM^ reverse transcriptase (50U/µl), 2 μL of a stem-loop RT primer, 1 μL of 5 X RT buffer, 0.13 μL of RNAse inhibitor (20U/µl), and 2.8 µl of nuclease-free water). We performed qPCR using 1 μL of cDNA with 7.60 µl of Taqman Universal PCR Master Mix (with No Amperase UNG, ABI) and 0.75 µl of miRNA-specific PCR TaqMan MicroRNA probes (20X, ABI) in a final reaction volume of 15 µl. All assays were run on an Applied Biosystems 7900HT system (ABI).

For TaqMan analysis mRNA expression, we used High-Capacity cDNA Reverse Transcription kit (ABI) with RNAses inhibitors to reversely transcribed RNA into cDNA. We followed the manufacturer’s instructions. Extracted total RNA was diluted at the initial concentration of 2 ng / μl. Briefly, we performed the reverse transcription reaction in a 20 μL total reaction volume containing 10 μL of extract RNA, 2 µl of RT buffer (10X), 0.8 µl of dNTPs (100 mM), 2 µl of RT random primers, 1 µl of MultiScribe^TM^ reverse transcriptase, 1µl of RNAse inhibitors and 3.2 µl of nuclease-free water. Among 6 candidates for endogenous controls (*Gapdh*, *Hmbs*, *RplpO*, *Ppia*, *B2M* and *Hprt1*), we selected *Hmbs* and *RplpO* by using GeNorm algorithm in qbase+ software. First, the optimal number of reference targets was 2 (geNorm V <0.15). Then, *RplpO* and *Hmbs* constituted reference targets with very high stability (average geNorm M ≤ 0.2). We then amplified the cDNA by real-time PCR using TaqMan Fast Advanced Master Mix (2X, ABI) and mRNA-specific TaqMan assays (20X, ABI, Thermofisher, see Supplementary Table 2) according to the manufacturer’s protocol. For all experiments, we ran triplicates for each reaction and we calculated the average Cq values. We included a no template control as negative control in all qRT-PCR reactions. Comparison between groups was made using the method of 2^−ΔΔCq^ with U87 and snoRNA as the endogenous controls for miRNA, *Hmbs* and *RplpO* for mRNAs.

### Immunoblotting

Tissue punches of DS (bilateral 2 mm^2^ punches obtained from a 2.1 mm coronal section using a brain matrix, bregma: from 1.7 to -0.40 mm) were collected at least 30 min after the end of the last session of heroin self-administration and stored at −80°C. Tissue was homogenized in ice-cold homogenization buffer (25 mM HEPES-NaOH pH 7.4, 0.1 % NP-40, 500 mM NaCl, 1 mM EDTA, 20 mM Dithiothreitol, 20 mM NaF, protease inhibitors cocktail (Roche)) using TissueRuptor (Qiagen) and then centrifuged (14 000 rpm for 2 minutes at 4°C, one sample containing the striata from both hemispheres of one individual). Supernatant was recovered and proteins were quantified with Bradford assay (Sigma Aldrich) and samples were stored at -80°C until further used. Proteins (30 µg/lane, a quantity determined with a calibration curve (data not shown)) were resolved on 8 % SDS-PAGE after heat denaturation (5 minutes, 95°C) in sample buffer (62.5 mM Tris-HCl, pH 6.8, 2% SDS, 10% glycerol, 5% β-mercaptoethanol, 0.001% bromophenol blue)^63^, transferred to PVDF membrane, and cut into 2 pieces (the top was used for Sez6 probing and the bottom for GAPDH detection).

The membranes were washed in Tris-buffered saline (20 mM Tris-HCl, pH 7.4, 137 mM NaCl) (TBS), incubated in blocking buffer (TBS/0.05 % Tween-20 (TBS/T)/5 % non-fat dried milk) for 1 hour at room temperature (RT), and then with the primary antibodies diluted in the blocking buffer (rabbit polyclonal anti-SEZ6^33^ diluted at 1:1000, mouse monoclonal anti-GAPDH (Thermofischer Scientific cat no. MA5-15738) diluted at 1:50,000) overnight at 4°C (for anti-SEZ6 antibody) or for one hour at RT (for anti-GAPDH antibody). After washes with TBS/T (4 x 10 min), membranes were incubated with secondary antibodies (from LI-COR Biosciences): anti-rabbit IRDye 800CW (cat #925-32211) and anti-mouse IRDye 680RD (cat #925-68070) diluted at 1:10 000 according to manufacturer’s instructions, for 1 hour at RT. After washes with TBS/T (6 x 10 min), antibodies were revealed with the Odyssey system (LI-COR Biosciences) and protein bands were quantified with Image Studio Lite (LI-COR Biosciences).

### In vitro dual luciferase assays

A part of the 3’-UTR of *Sez6* (ACTGGCGCACATTACCAGAAGCCAACATTCTGCTTCTCTTTT**gccctgga**CCTCCA**gccctg ga**AGTTGCAGGCAGAGAAGGGGCACGTCTTACACATAGCCACCATCCA**gccctgg**CATGCAAGGACTCCAGAAGAGCTG) containing predicted targeted sequences (in lower case) of miR-3594-5p was amplified from rat brain cDNA and cloned downstream of the Renilla luciferase reporter gene into the psiCHECK-2 vector (Promega).

As a control, another region of the 3’-UTR of *Sez6* (ACTCTATCCCAAGCTCAGGTTGGGCCTCACTGGATACCCATGCTCCACACCAAAGCAG GACACCCTATGGCTCCCTGGCATCCCTCTACCCTGAGGAGGGAAAGCCCTTCCTTGAAATTATCTGGCTGTTACAAATGTCC) lacking target sequences of miR-3594-5p was cloned in psiCHECK-2. Recombinant vectors were transfected in HEK293 cells with 10 nM of miRNA mimics (miR-3594-5p, mature sequence: CCCAGGGCAGAGCAGUGUGAA; miR-negative control, ThermoFisher scientific) with Lipofectamine 2000 (Invitrogen). After 48 hours, a dual-luciferase reporter assay (Promega) was performed to measure Firefly and Renilla luciferase activity. Renilla activity was normalized to the Firefly luciferase activity. All the experiments were performed in triplicate and repeated at least three times.

### Drugs

Heroin (3,6-diacetylmorphine HCl, Francopia, Anthony, France) and Cocaine (Cocaine HCl, Cooper-Melun, France) were dissolved in a sterile physiological saline solution (0.9 % w/v, B. Braun) in 250-ml sterile bags at concentrations of 0.20 mg / ml and 3.38 mg / ml, respectively. We used unit doses for both drugs based on our previous studies^29, 52, 53, 56^.

### Statistical analyses

We analyzed the data with the scientific graphing and statistical software GraphPad Prism version 7.00 (San Diego, USA). We checked normal distribution and used parametric tests when this assumption was confirmed. Data were tested for normality using the Shapiro-Wilk normality tests. Homogeneity of variance between groups was tested using the Brown and Forsythe’s test. For the first hour of the self-administration sessions, numbers of injections and lever presses (active and inactive) were subjected to two-way mixed analyses of variance (ANOVA) with a between-subjects factor (experimental groups: ShA versus LgA versus LgA-Yoked groups) and a within-subjects factor (self-administration sessions). For the 5 remaining hours in LgA group, data were analyzed with one-way repeated measures ANOVA. We applied the Greenhouse-Geisser correction for repeated-measures ANOVA when the assumption of sphericity was violated. All post hoc comparisons for significant interactions were carried out by the Bonferonni post hoc test. Results were considered statistically significant with a value of p<0.05. The comparisons between experimental groups of miRNA and mRNA expression were done on values normalized to the Naive group and expressed relative to the endogenous control levels by using the method of 2^−ΔΔCq^. Only Cq values having a standard deviation of the triplicate above 0.5 were excluded. After testing the assumption of normality and homogeneity of variance, fold-change data were analyzed with one-way ANOVA, followed by a Bonferroni’s post hoc test. Without normal distribution, we used the non-parametric test of Kruskal-Wallis followed by a post hoc Dunn test. Outliers were detected by using the ROUT’s test (Robust regression and Outlier removal, Q = 1%) and excluded. During the behavioral experiments, the exclusion of one pilot LgA rat led automatically to the exclusion of the matched LgA-yoked rat. In Experiment 6, Pearson’s correlation coefficients were calculated between individual slope coefficients of escalation in heroin intake during the first hour of self-administration and *Sez6* mRNA levels expressed within the dorsal striatum. Since heroin intake increased essentially from session 4 to 14, individual slope coefficients were computed fitting the data over this window time period (Fig. 4d). In addition, total cumulative heroin intake (per mg/kg of body weight), obtained during the whole duration of the Experiment 6, was correlated to the *Sez6* mRNA expression. Supplementary Table 3 provides a report of the statistical analyses for the results presented in the figures.

## Acknowledgements

This work was supported by the Centre National de la Recherche Scientifique CNRS UMR ERL-3649 and the French National Agency (grant ANR-12-SAMA-0003). Behavioral studies were performed at the animal core facility of BioMedTech Facilities and molecular biology data were generated at the molecular biology core facility of BioMedTech Facilities, INSERM US36 | CNRS UMS2009 | Université de Paris. I.B. was supported by an international doctoral fellowship from Sorbonne Paris Cité, M.B. by a LABEX PRINCEPS funding and L.C. by a doctoral fellowship from CNRS. Addiction research in J.M.G.’s lab is supported by an Australian National Health and Medical Research Council Grant (GNT1140050). We thank Benjamin Saintpierre for generating heatmaps, Drs. Jacques Pantel and Hélène Geoffroy for fruitful discussions and Drs Michel Engeln and Karine Guillem for their helpful comments on a previous version of the manuscript.

## Author contributions

M.L., F.N., N.M. and S.H.A. conceived the project and designed the experiments. M.L., I.B., M.B., and C.C. performed the behavioral experiments. S.J. and A.D. performed the microarray assays and contributed to bioinformatics analyses. M.L., I.B., I.N., M.B., S.M-L, S. M-R conducted microRNA and RNA extraction and quantification. J.M.G provided a reagent and contributed to methodology. L.C., I.N. and N.M. performed in vitro luciferase assays and immunoblot analysis. M.L., I.B., L.C., I.N., M.B., N.M. analyzed the data. M.L., F.N., N.M. and S.H.A. wrote the paper with contributions from all authors.

## Ethics Declarations

The authors declare no competing interests.

## Supplementary information

**Extended Data Fig.1:**
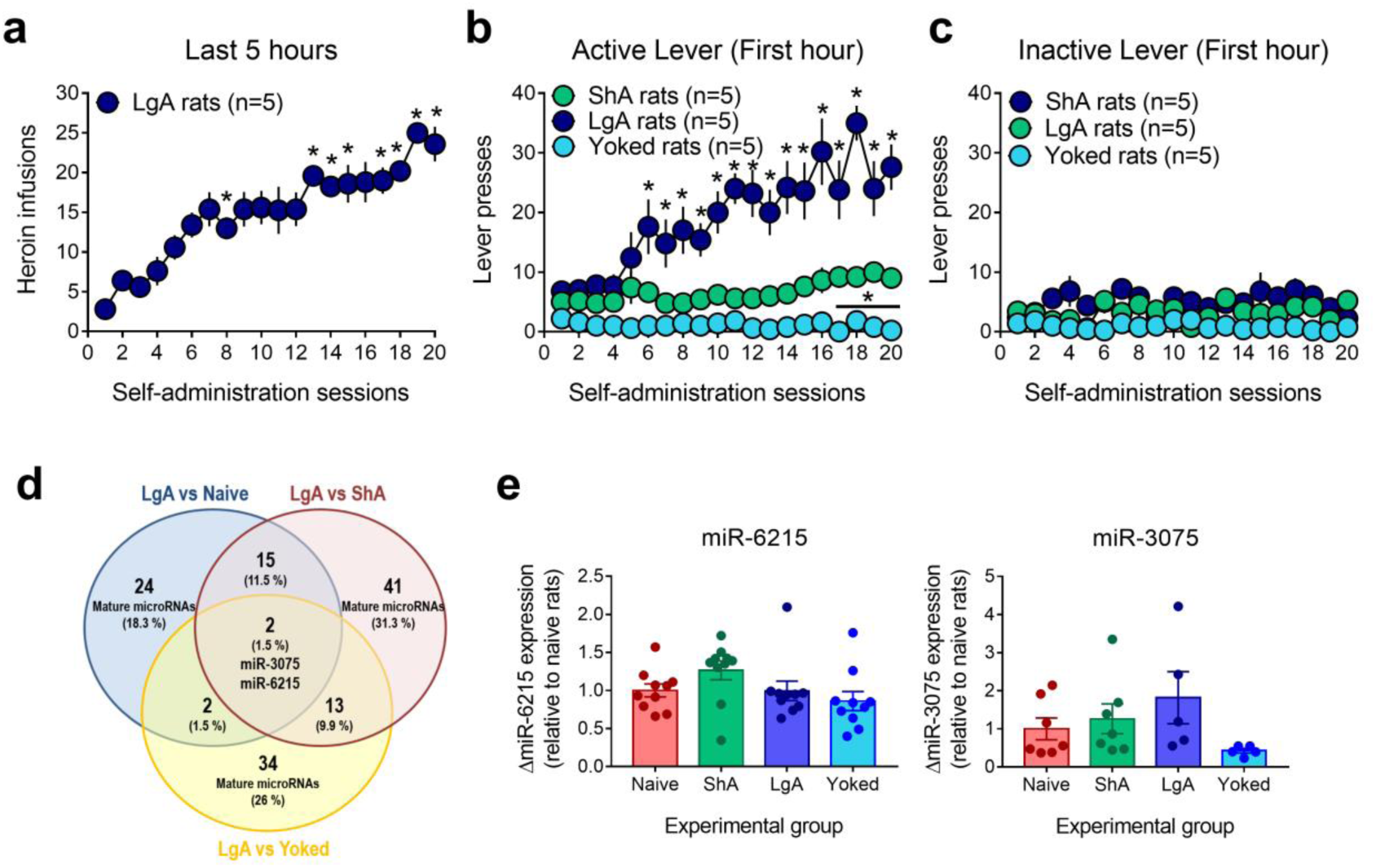
miRNA expression profiling in rat dorsal striatum after 20 days of differential access to heroin using miRNA microarrays. **a)** Heroin infusions earned during the 5 remaining hours of heroin self-administration over 20 daily sessions in extended access rats. *, different from the first session, *P<0.05; Bonferroni’s post hoc test. **b)** Active-lever and **c)** inactive-lever presses (mean ± s.e.m.) during the first hour of heroin self-administration over 20 daily sessions in restricted (ShA rats, green circles, 1h/day, n=5), extended access (LgA rats, dark blue circles, 6h/day, n=5) and heroin-yoked rats (Yoked rats, light blue circles, 6h/day, n=5). *, different from ShA rats, *P<0.05, Bonferroni’s multiple-comparisons test). **d)** Venn diagram representing the overlap of mature miRNAs significantly differentially expressed in the dorsal striatum according to the drug history (n=5 rats per group). MiR-3075 and miR-6215 were overexpressed only in LgA rats compared with the other groups. **e)** Taqman qRT-PCR analyses of microRNA expression for miR-6215 (n=10 rats per group) and miR-3075 (n= 5-7 rats per group) in LgA rats compared with heroin-naive, ShA and Yoked rats. For qRT-PCR analyses, relative microRNA expression levels were calculated using the method of 2−ΔΔCt with snoRNA and U87 as the endogenous controls for normalization. Statistical details are included in Supplementary Table 3.

**Extended Data Fig. 2:**
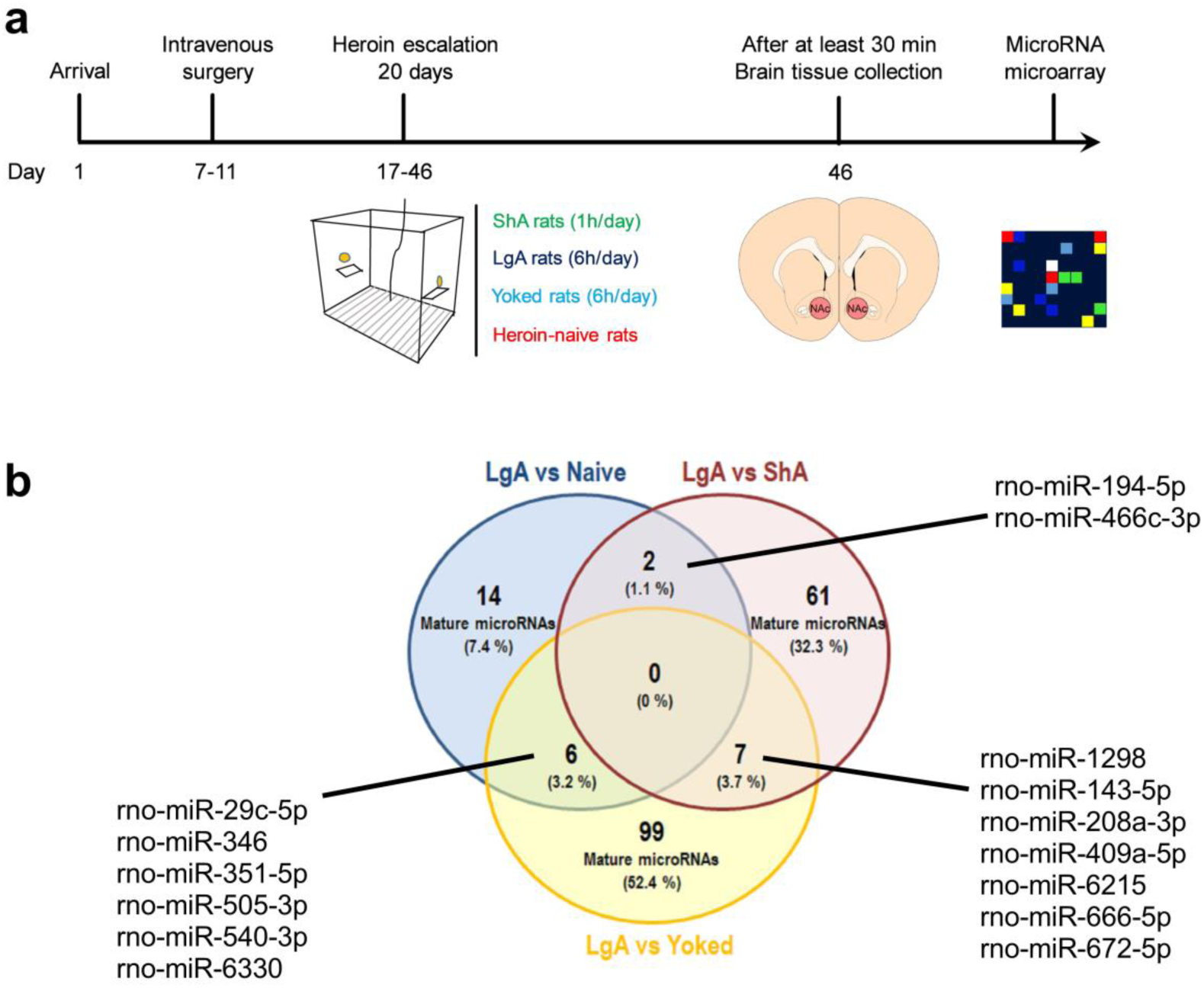
miRNA expression profiling in rat nucleus accumbens after 20 days of differential access to heroin using miRNA microarrays. **a)** Timeline of the experimental procedure. **b)** Venn diagram representing the overlap of mature miRNAs significantly differentially expressed in the nucleus accumbens according to the drug history (n=5 rats per group). No miRNA was altered specifically in LgA rats compared with the 3 other experimental groups.

**Extended Data Fig. 3:**
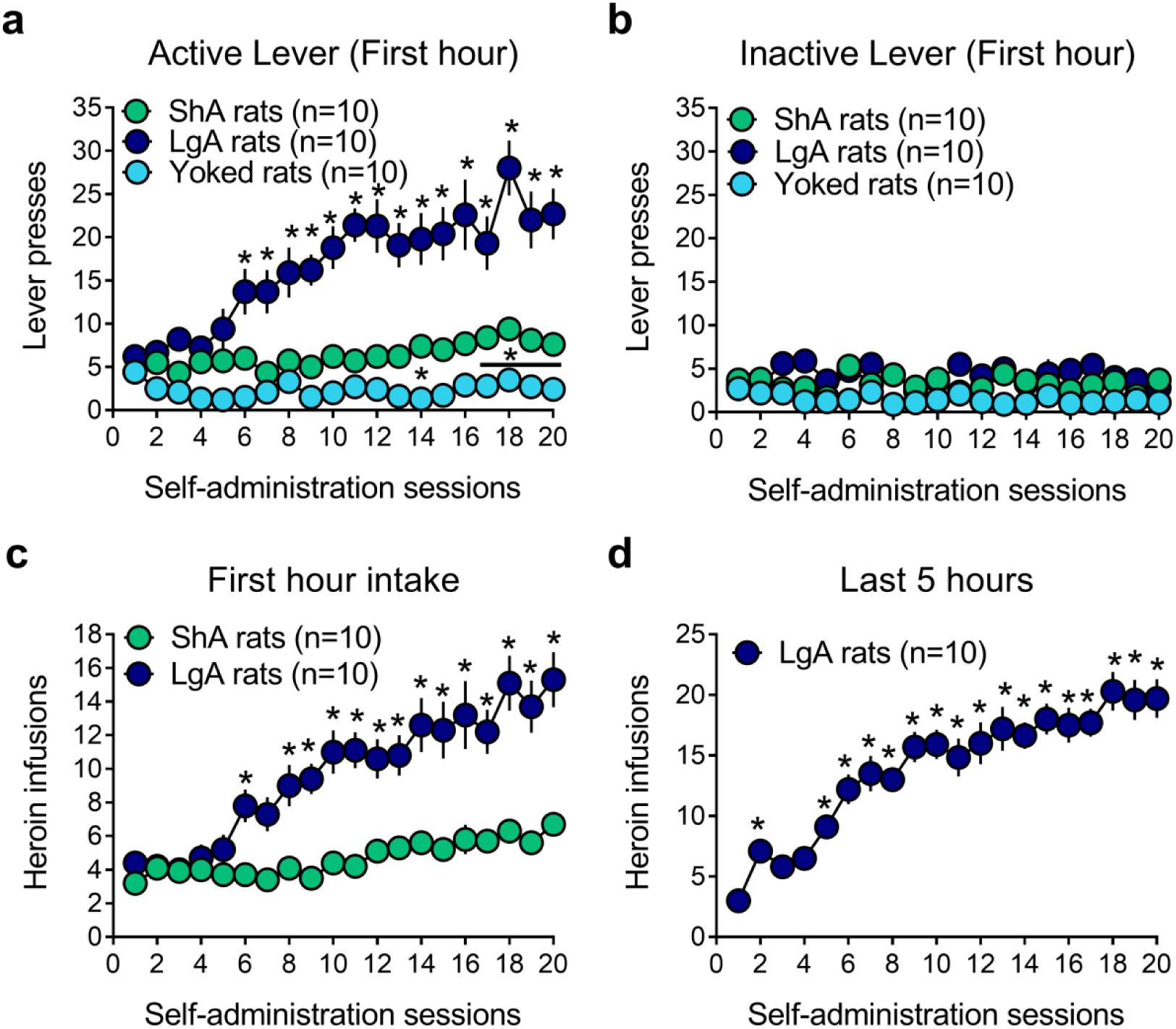
Escalation of heroin intake in extended access rats to measure miRNA expression in the dorsal striatum by qRT-PCR. **a)** Active-lever and **b)** inactive-lever presses (mean ± s.e.m.) during the first hour of heroin self-administration over 20 daily sessions in restricted (ShA rats, green circles, 1h/day, n=10), extended access (LgA rats, dark blue circles, 6h/day, n=10) and heroin-yoked rats (Yoked rats, light blue circles, 6h/day, n=10). *, different from ShA rats, *P<0.05, Bonferroni’s multiple-comparisons test. **c)** Heroin infusions earned during the first hour of heroin self-administration over 20 daily sessions in restricted and extended access rats. Post hoc comparisons showed that significant differences between groups appeared from session 6 onward (*different from ShA rats, P<0.05, Bonferroni’s multiple-comparisons test). **d)** Heroin infusions earned during the 5 remaining hours of heroin self-administration over 20 daily sessions in extended access rats. *, different from the first session, *P<0.05; Bonferroni’s post hoc test. Data are presented as mean ± s.e.m. N=10 rats / experimental groups. Statistical details are included in Supplementary Table 3.

**Extended Data Fig.4:**
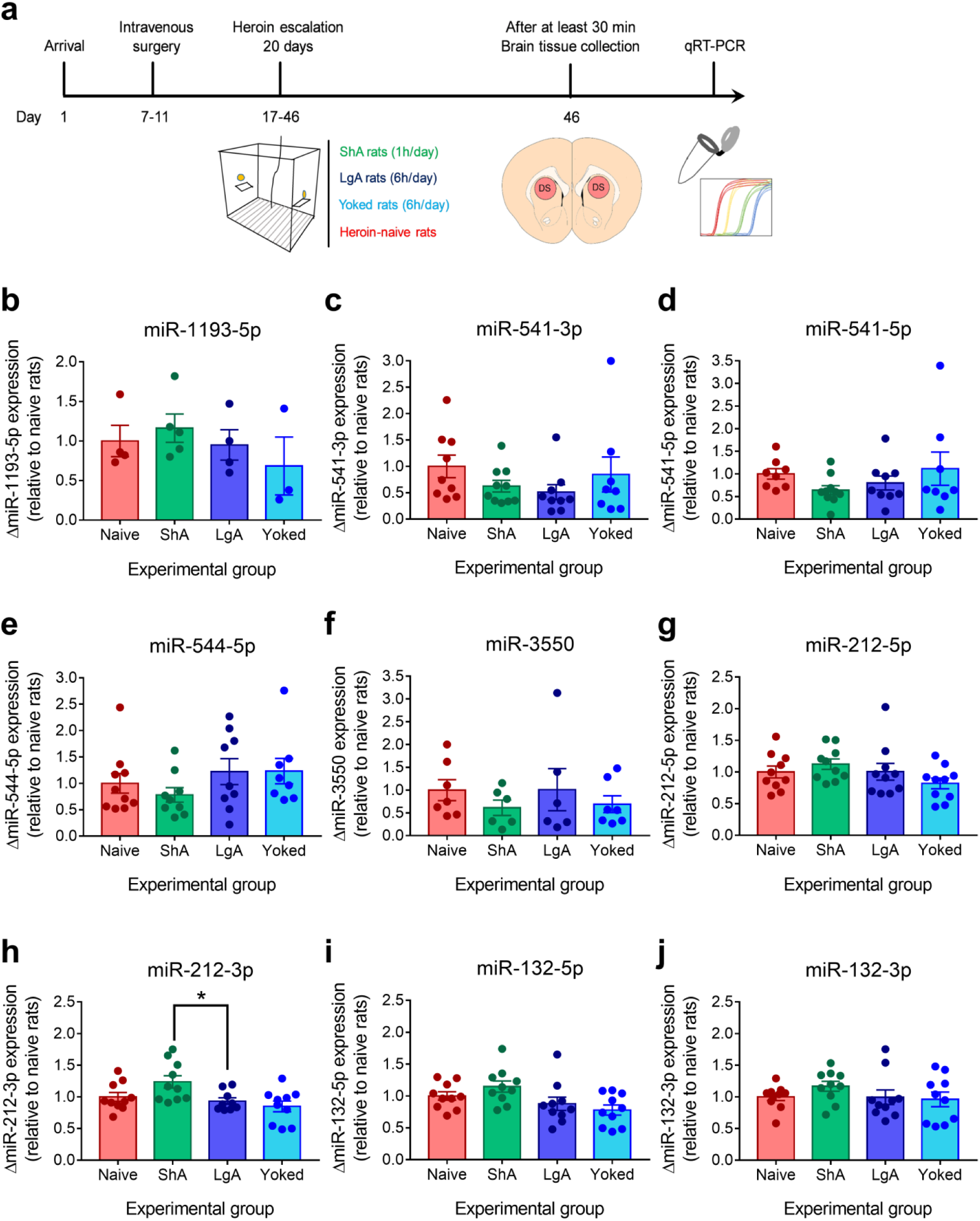
Expression of candidate miRNAs in the dorsal striatum measured by Taqman qRT-PCR following 20 days of differential access to heroin. **a)** Timeline of the experimental procedure. **b-j)** Taqman qRT-PCR did not validate alterations specific to LgA rats in miR-1193-5p, miR-541-3p, miR-541-5p, miR-544-5p and miR-3550 levels. No alteration in the expression levels of miR-212-5p, miR-212-3p, miR-132-5p and miR-132-3p was also confirmed by qRT-PCR assays (one-way ANOVA or Kruskal-Wallis ANOVA, n=3-10 per group). *different from LgA rats, P<0.05; Dunn’s multiple comparison test. Data are presented as mean ± s.e.m. For qRT-PCR analyses, relative microRNA expression levels were calculated using the method of 2−ΔΔCt with snoRNA and U87 as the endogenous controls for normalization. Statistical details are included in Supplementary Table 3.

**Extended Data Fig.5:**
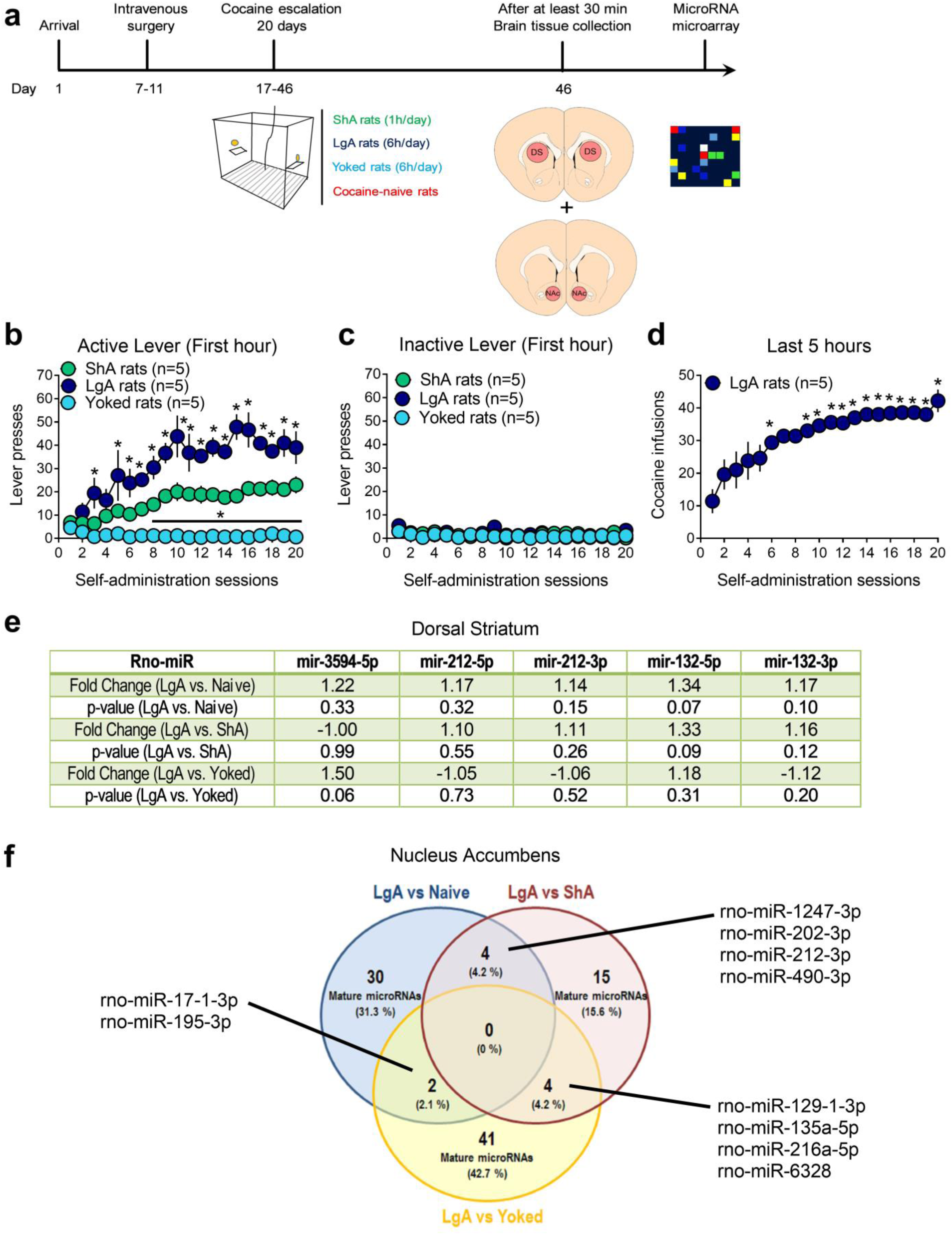
miRNA expression analysis in rat nucleus accumbens and dorsal striatum after 20 days of differential access to cocaine by microarray. **a)** Timeline of the experimental procedure. **b)** Active-lever and **c)** inactive-lever presses (mean ± s.e.m.) during the first hour of cocaine self-administration over 20 daily sessions in restricted (ShA rats, green circles, 1h/day, n=5), extended access (LgA rats, dark blue circles, 6h/day, n=5) and cocaine-yoked rats (Yoked rats, light blue circles, 6h/day, n=5). *, different from ShA rats, *P<0.05, Bonferroni’s multiple-comparisons test. **d)** Cocaine infusions earned during the 5 remaining hours of cocaine self-administration over 20 daily sessions in extended access rats. *, different from the first session, *P<0.05; Bonferroni’s post hoc test. Data are presented as mean ± s.e.m. **e)** Table summarizing expression levels of miR-3594-5p, miR-212 and miR-132 from the same cluster, measured by microarray analysis. No alteration in the DS of LgA rats was found. **f)** Venn diagram representing the overlap of mature miRNAs significantly differentially expressed in the nucleus accumbens according to the cocaine history (n=5 rats per group). No miRNA was altered specifically in LgA rats compared with the 3 other experimental groups. N=5 rats / experimental groups. Statistical details are included in Supplementary Table 3.

**Extended Data Fig.6:**
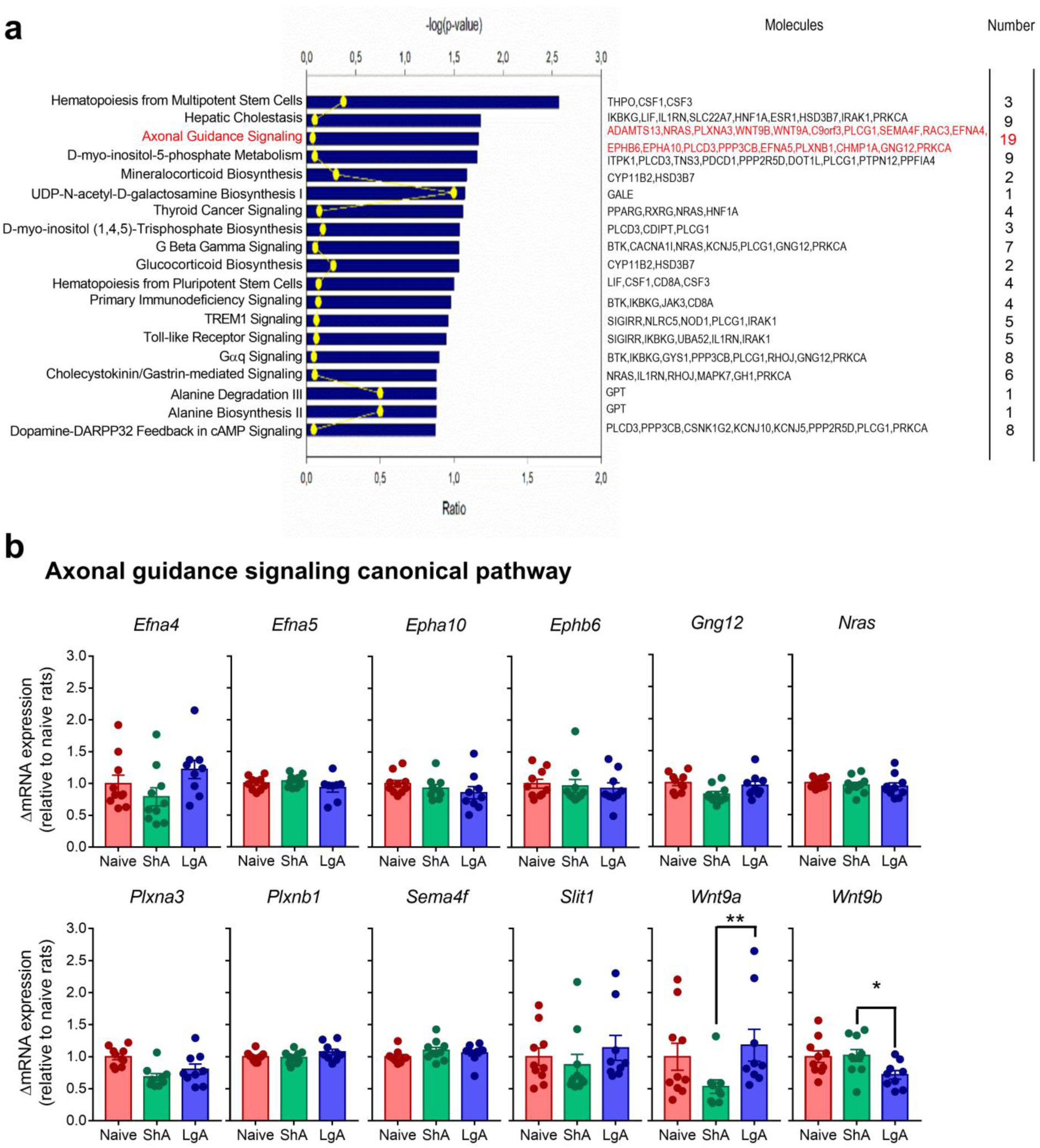
Top canonical pathways enriched by predicted targets of miR-3594-5p. **a)** Top 19 significantly enriched IPA canonical pathways of non-experimentally validated target genes of miR-3594-5p. Note that axonal guidance signaling pathway comprise 19 putative targets of miR-3594-5p. **b)** No significant difference in the relative dorsal striatal mRNA levels of 12 target genes contributing to the axon guidance signaling pathway in LgA (n=10) compared with heroin-naïve (n=10) and ShA (n=10) rats. Data are presented as mean ± s.e.m. For qRT-PCR analyses, relative mRNA expression levels were calculated using the method of 2−ΔΔCt with *Rplpo* and *Hmbs* as the endogenous controls for normalization. Statistical details are included in Supplementary Table 3.

**Extended Data Fig.7:**
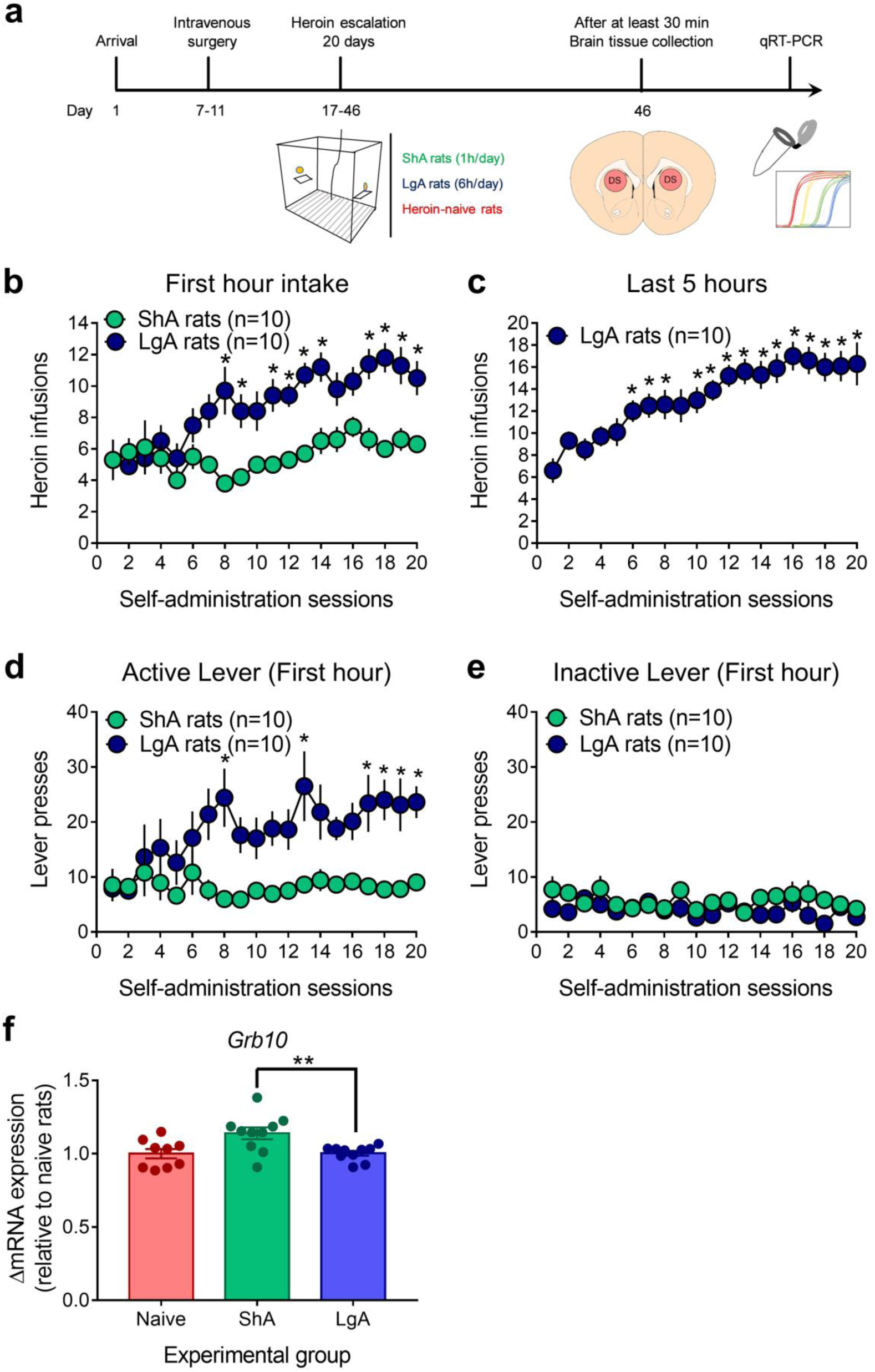
Target mRNA expression levels determined by qRT-PCR in rats after heroin escalation. **a)** Timeline of the experimental procedure. **b)** Heroin infusions earned during the first hour of heroin self-administration over 20 daily sessions in restricted (ShA rats, green circles, 1h/day, n=10) and extended access rats (LgA rats, dark blue circles, 6h/day, n=10). *different from ShA rats, P<0.05, Bonferroni’s multiple-comparisons test. **c)** Heroin infusions earned during the 5 remaining hours of heroin self-administration over 20 daily sessions in extended access rats. *, different from the first session, *P<0.05; Bonferroni’s post hoc test. **d)** Active-lever and **e)** inactive-lever presses during the first hour of heroin self-administration over 20 daily sessions in restricted and extended access. *, different from ShA rats, *P<0.05, Bonferroni’s multiple-comparisons test. **f)** Taqman qRT-PCR analysis of mRNA expression for *Grb10* in LgA (n=10) rats compared with heroin-naïve (n=9) and ShA (n=10) rats. Data are presented as mean ± s.e.m. For qRT-PCR analyses, relative mRNA expression levels were calculated using the method of 2−ΔΔCt with *Rplpo* and *Hmbs* as the endogenous controls for normalization. Statistical details are included in Supplementary Table 3.

**Extended Data Fig.8:**
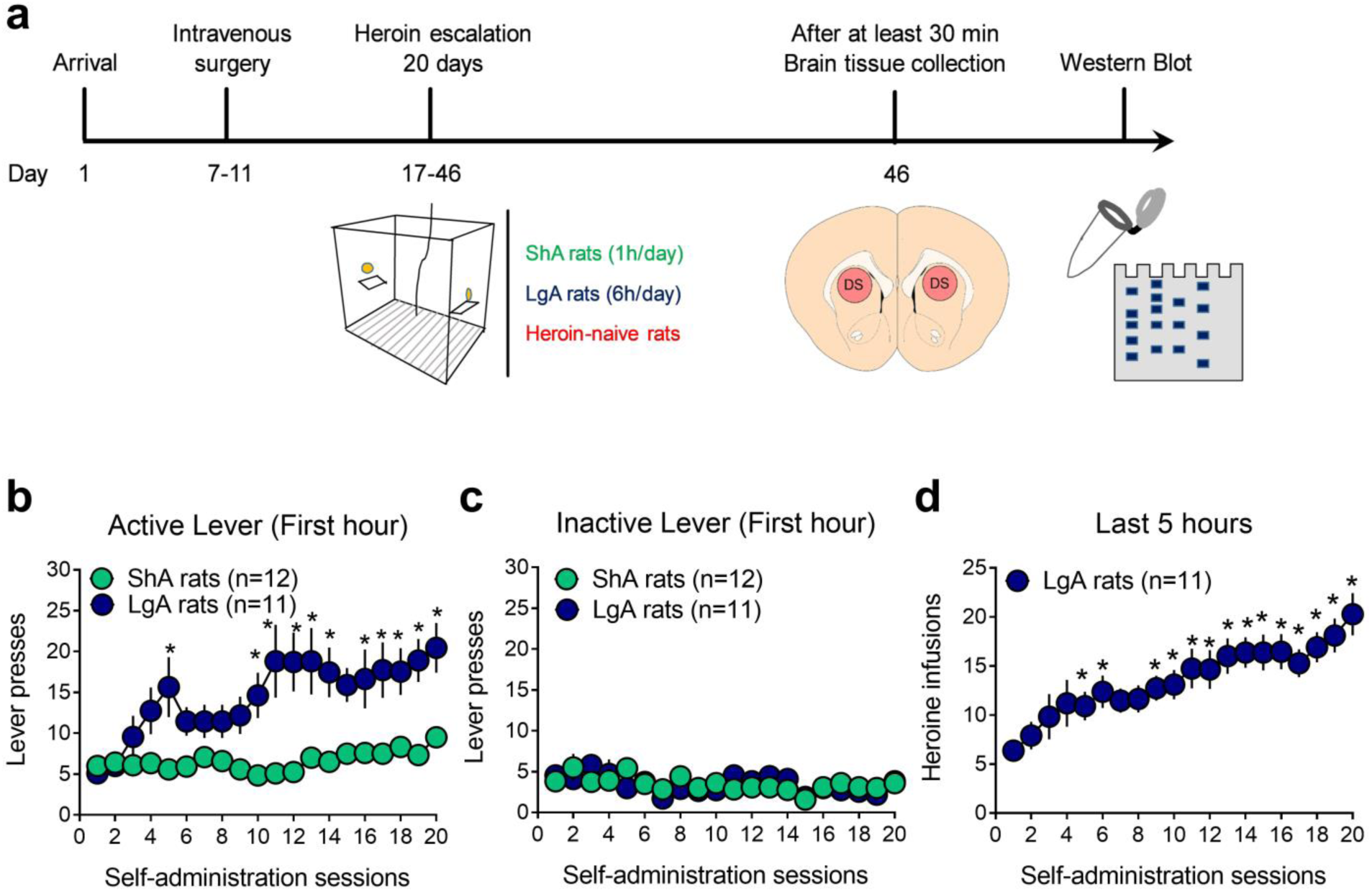
Mature SEZ6 protein is inhibited in the dorsal striatum of extended access rats following 20 days of heroin self-administration. **a)** Timeline of the experimental procedure. **b)** Active-lever and **c)** inactive-lever presses during the first hour of heroin self-administration over 20 daily sessions in restricted (ShA rats, green circles, 1h/day, n=12) and (LgA rats, dark blue circles, 6h/day, n=11) extended access rats. *, different from ShA rats, *P<0.05, Bonferroni’s multiple-comparisons test. **d)** Heroin infusions earned during the 5 remaining hours of heroin self-administration over 20 daily sessions in extended access rats. *, different from the first session, *P<0.05; Bonferroni’s post hoc test. Data are presented as mean ± s.e.m. Statistical details are included in Supplementary Table 3.

**Supplementary Table 1:** Primers for miRNA qRT-PCR analysis.

MicroRNA assays
| Mature miRNA | miRBase Accession Number (v22) | Taqman probe (Assay ID) |
| --- | --- | --- |
| rno-miR-6215 | MIMAT0024854 | 473573_mat |
| rno-miR-3075 | MIMAT0025057 | 476103_mat |
| rno-miR-27a-5p | MIMAT0004715 | 002445 |
| rno-miR-3594-5p | MIMAT0017898 | 462624_mat |
| rno-miR-1193-5p | MIMAT0017858 | 462317_mat |
| rno-miR-541-3p | MIMAT0017213 | 461954_mat |
| rno-miR-541-5p | MIMAT0003177 | 002562 |
| rno-miR-544-5p | MIMAT0017359 | 464784_mat |
| rno-miR-3550 | MIMAT0017808 | 462851_mat |
| rno-miR-212-5p | MIMAT0017158 | 463930_mat |
| rno-miR-212-3p | MIMAT0000883 | 002551 |
| rno-miR-132-5p | MIMAT0017123 | 002132 |
| rno-miR-132-3p | MIMAT0000838 | 000457 |
| SnoRNA | U64702 | 001718 |
| U87 | AF272707 | 001712 |
| 4.5S RNA(H) | AY228147 | 001716 |
| U6 | NR_004394 | 001973 |

**Supplementary Table 2:** Primers for mRNA qRT-PCR analysis.

| **Gene ID** | **Description** | **Taqman probe (Assay ID)** |
| --- | --- | --- |
| *Efna4* | Ephrin A4 | Rn01455418_m1 |
| *Efna5* | Ephrin A5 | Rn01427154_m1 |
| *Epha10* | Ephrin type-A receptor 10 | Rn01519668_m1 |
| *Ephb6* | Ephrin type-B receptor 6 | Rn01456013_g1 |
| *Gng12* | G protein subunit gamma 12 | Rn01425123_m1 |
| *Nras* | NRAS proto-oncogene, GTPase | Rn00821645_g1 |
| *Plxna3* | Plexin A3 | Rn01529862_m1 |
| *Plxnb1* | Plexin B1 | Rn01746615_m1 |
| *Sema4f* | Semaphorin 4F | Rn00570562_m1 |
| *Slit1* | Slit guidance ligand 1 | Rn00576227_m1 |
| *Wnt9a* | Wnt family member 9A | Rn01496604_m1 |
| *Wnt9b* | Wnt family member 9B | Rn00457102_m1 |
| *Grb10* | Growth factor receptor bound protein 10 | Rn01401196_m1 |
| *Sez6* | Seizure related 6 homolog | Rn01751445_m1 |
| *Gapdh* | Glyceraldehyde-3-phosphate dehydrogenase | Rn99999916_s1 |
| *Hmbs* | Hydroxymethylbilane synthase | Rn00565886-m1 |
| *Rplpo* | Ribosomal protein, large, P0 | Rn00821065_g1 |
| *Ppia* | Peptidylprolyl isomerase A | Rn00690933-m1 |
| *B2m* | Beta-2 microglobulin | Rn00560865-m1 |
| *Hprt1* | Hypoxanthine phosphoribosyltransferase 1 | Rn01527840-m1 |

**Supplementary Table 3:**
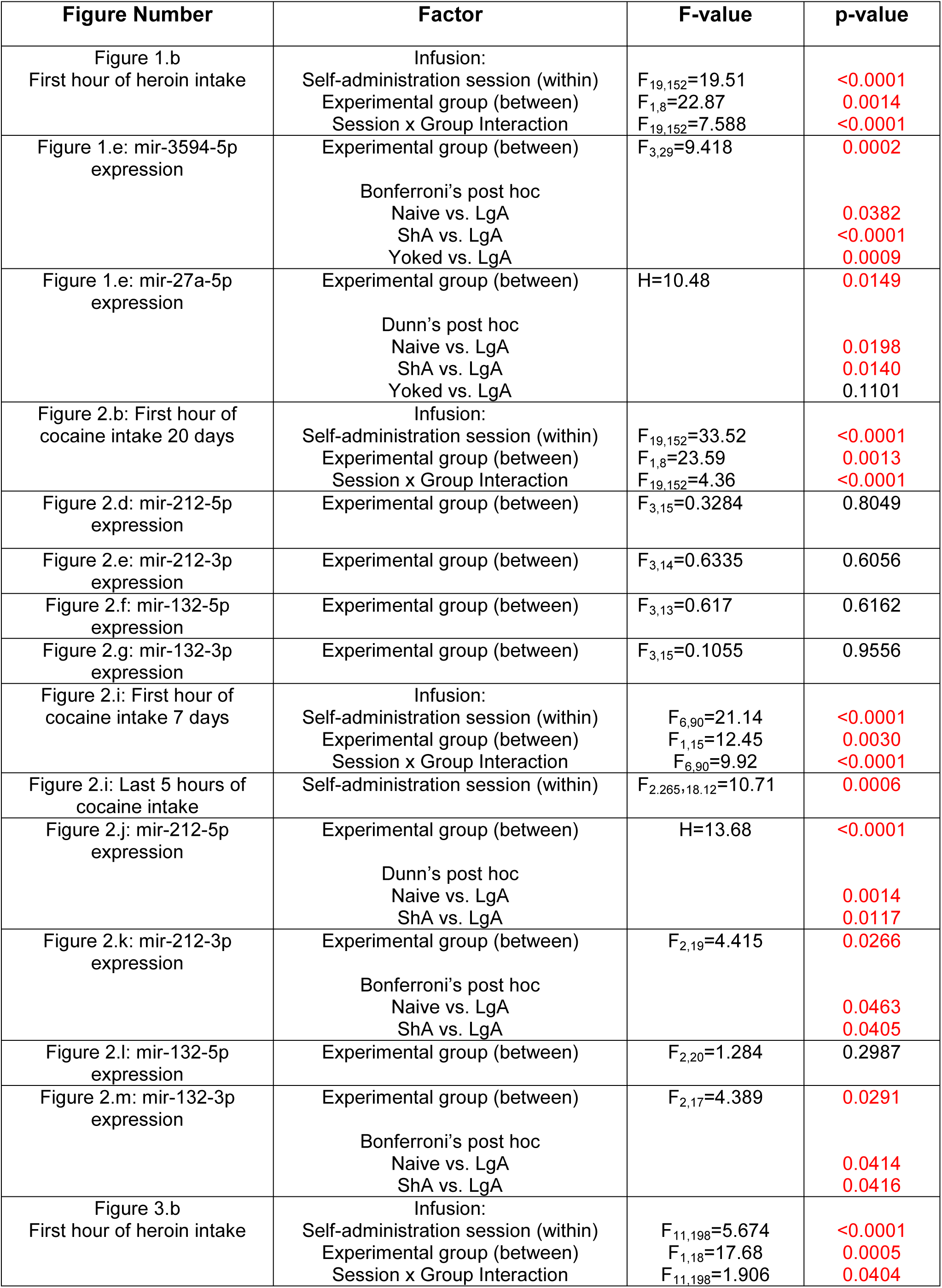
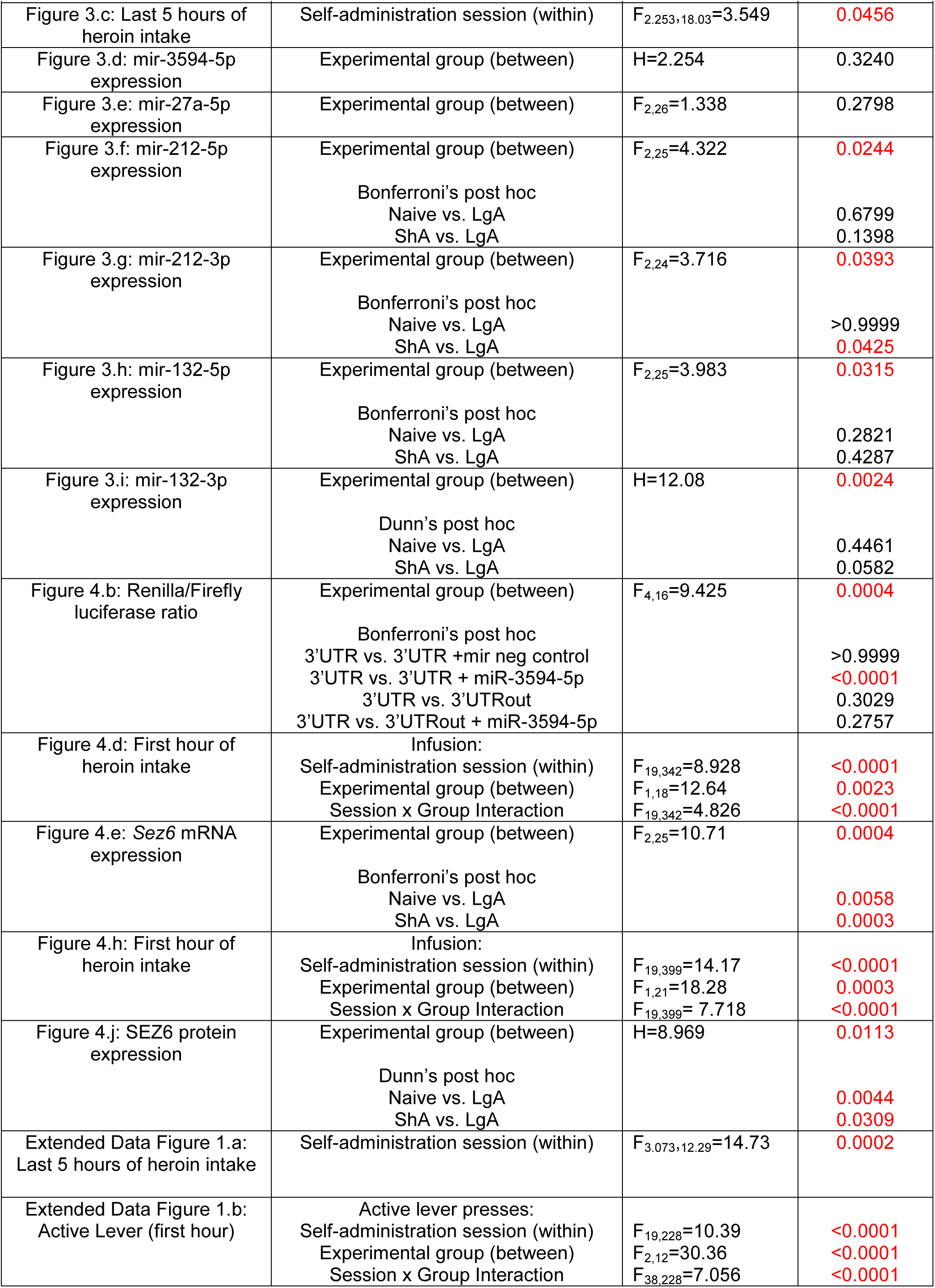
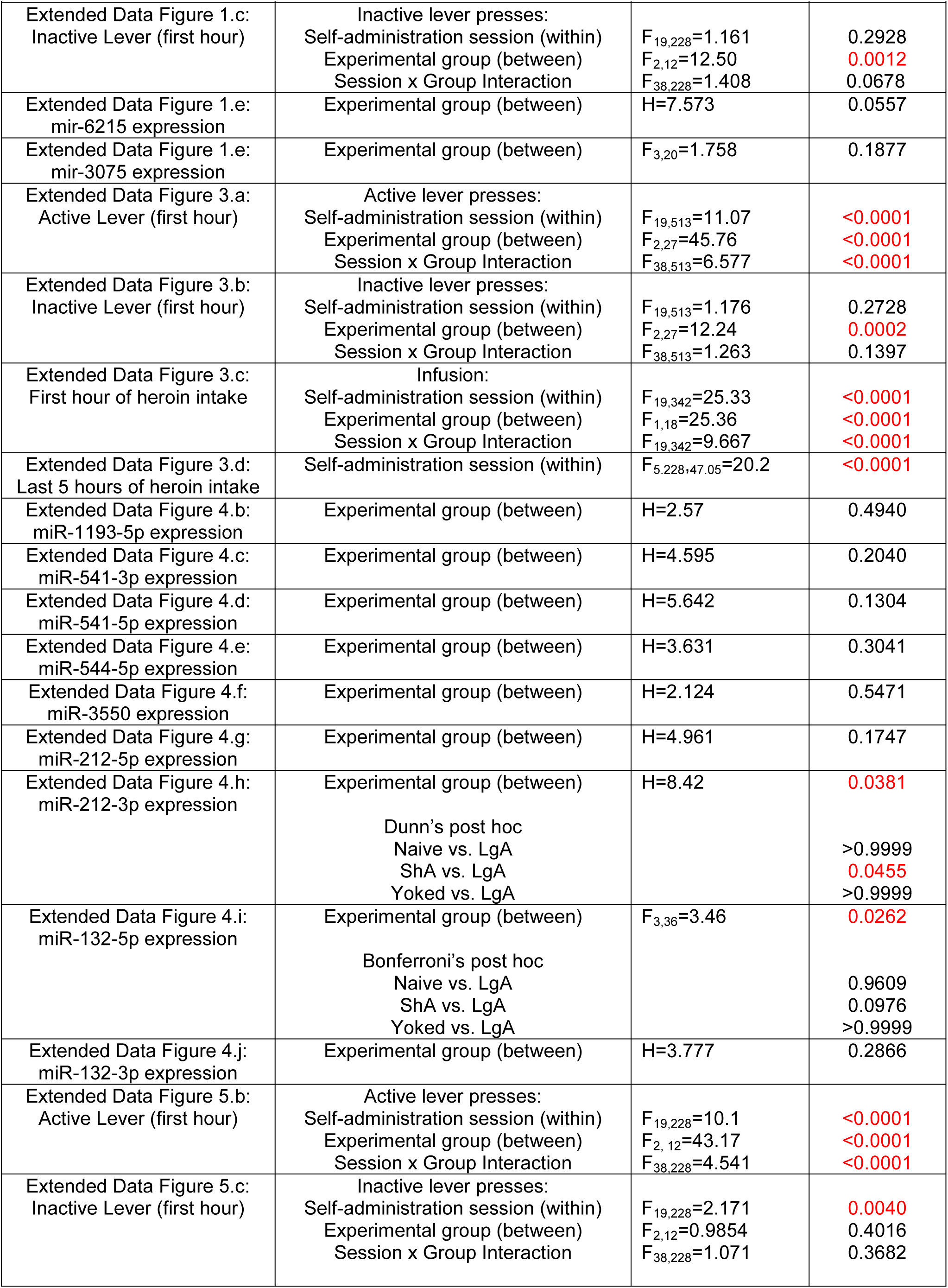
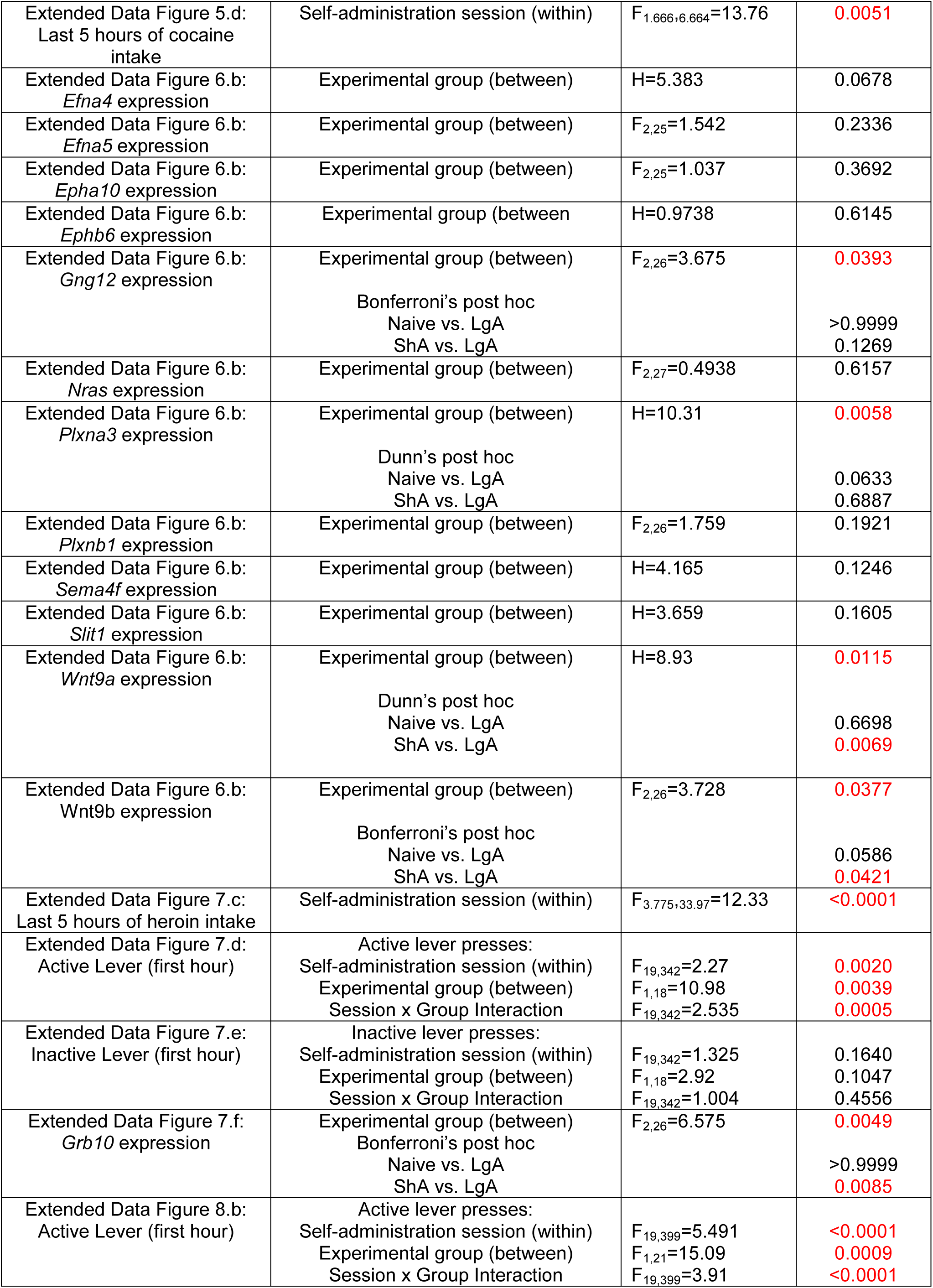
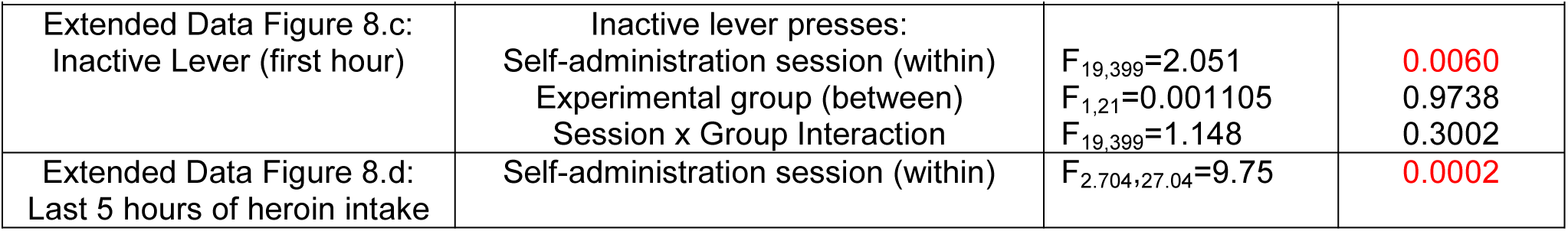
Statistical analysis.

## REFERENCES

1. Ahmed, S.H. & Koob, G.F. Transition from moderate to excessive drug intake: change in hedonic set point. Science 282, 298–300 (1998).

2. Vanderschuren, L. & Ahmed, S.H. Animal Models of the Behavioral Symptoms of Substance Use Disorders. Cold Spring Harbor perspectives in medicine (2020).

3. Kalivas, P.W. & O’Brien, C. Drug addiction as a pathology of staged neuroplasticity. Neuropsychopharmacology : official publication of the American College of Neuropsychopharmacology 33, 166–180 (2008).

4. Kasanetz, F., et al. Transition to addiction is associated with a persistent impairment in synaptic plasticity. Science 328, 1709–1712 (2010).

5. Kauer, J.A. & Malenka, R.C. Synaptic plasticity and addiction. Nature reviews. Neuroscience 8, 844–858 (2007).

6. Brown, A.L., Flynn, J.R., Smith, D.W. & Dayas, C.V. Down-regulated striatal gene expression for synaptic plasticity-associated proteins in addiction and relapse vulnerable animals. The international journal of neuropsychopharmacology 14, 1099–1110 (2011).

7. Vanderschuren, L.J., Di Ciano, P. & Everitt, B.J. Involvement of the dorsal striatum in cue-controlled cocaine seeking. The Journal of neuroscience : the official journal of the Society for Neuroscience 25, 8665–8670 (2005).

8. Belin, D. & Everitt, B.J. Cocaine seeking habits depend upon dopamine-dependent serial connectivity linking the ventral with the dorsal striatum. Neuron 57, 432–441 (2008).

9. Hodebourg, R., et al. Heroin seeking becomes dependent on dorsal striatal dopaminergic mechanisms and can be decreased by N-acetylcysteine. The European journal of neuroscience 50, 2036–2044 (2019).

10. Ambros, V. The functions of animal microRNAs. Nature 431, 350–355 (2004).

11. Bartel, D.P. MicroRNAs: genomics, biogenesis, mechanism, and function. Cell 116, 281–297 (2004).

12. Dreyer, J.L. New insights into the roles of microRNAs in drug addiction and neuroplasticity. Genome medicine 2, 92 (2010).

13. Ahmed, S.H. & Kenny, P.J. Cracking the molecular code of cocaine addiction. ILAR journal 52, 309–320 (2011).

14. Im, H.I. & Kenny, P.J. MicroRNAs in neuronal function and dysfunction. Trends in neurosciences 35, 325–334 (2012).

15. Bali, P. & Kenny, P.J. MicroRNAs and Drug Addiction. Frontiers in genetics 4, 43 (2013).

16. Jonkman, S. & Kenny, P.J. Molecular, cellular, and structural mechanisms of cocaine addiction: a key role for microRNAs. Neuropsychopharmacology : official publication of the American College of Neuropsychopharmacology 38, 198–211 (2013).

17. Smith, A.C.W. & Kenny, P.J. MicroRNAs regulate synaptic plasticity underlying drug addiction. Genes, brain, and behavior 17, e12424 (2018).

18. Hollander, J.A., et al. Striatal microRNA controls cocaine intake through CREB signalling. Nature 466, 197–202 (2010).

19. Im, H.I., Hollander, J.A., Bali, P. & Kenny, P.J. MeCP2 controls BDNF expression and cocaine intake through homeostatic interactions with microRNA-212. Nature neuroscience 13, 1120–1127 (2010).

20. Tapocik, J.D., et al. Neuroplasticity, axonal guidance and micro-RNA genes are associated with morphine self-administration behavior. Addiction biology 18, 480–495 (2013).

21. Yan, B., et al. MiR-218 targets MeCP2 and inhibits heroin seeking behavior. Scientific reports 7, 40413 (2017).

22. Kim, J., Im, H.I. & Moon, C. Intravenous morphine self-administration alters accumbal microRNA profiles in the mouse brain. Neural regeneration research 13, 77–85 (2018).

23. Mavrikaki, M., et al. Overexpression of miR-9 in the Nucleus Accumbens Increases Oxycodone Self-Administration. The international journal of neuropsychopharmacology 22, 383–393 (2019).

24. Hsu, C.W., Huang, T.L. & Tsai, M.C. Decreased Level of Blood MicroRNA-133b in Men with Opioid Use Disorder on Methadone Maintenance Therapy. Journal of clinical medicine 8 (2019).

25. Gu, W.J., et al. Altered serum microRNA expression profile in subjects with heroin and methamphetamine use disorder. Biomedicine & pharmacotherapy = Biomedecine & pharmacotherapie 125, 109918 (2020).

26. Xu, W., et al. Increased expression of plasma hsa-miR-181a in male patients with heroin addiction use disorder. Journal of clinical laboratory analysis 34, e23486 (2020).

27. Xu, W., et al. Role of nucleus accumbens microRNA-181a and MeCP2 in incubation of heroin craving in male rats. Psychopharmacology (2021).

28. Ahmed, S.H., Walker, J.R. & Koob, G.F. Persistent increase in the motivation to take heroin in rats with a history of drug escalation. Neuropsychopharmacology : official publication of the American College of Neuropsychopharmacology 22, 413–421 (2000).

29. Lenoir, M. & Ahmed, S.H. Heroin-induced reinstatement is specific to compulsive heroin use and dissociable from heroin reward and sensitization. Neuropsychopharmacology : official publication of the American College of Neuropsychopharmacology 32, 616–624 (2007).

30. Ahmed, S.H. Escalation of drug use. Animal models of drug addiction. Neuromethods 53, 267–292 (2011).

31. Tapocik, J.D., et al. MicroRNAs Are Involved in the Development of Morphine-Induced Analgesic Tolerance and Regulate Functionally Relevant Changes in Serpini1. Frontiers in molecular neuroscience 9, 20 (2016).

32. Osaki, G., Mitsui, S. & Yuri, K. The distribution of the seizure-related gene 6 (Sez-6) protein during postnatal development of the mouse forebrain suggests multiple functions for this protein: an analysis using a new antibody. Brain research 1386, 58–69 (2011).

33. Gunnersen, J.M., et al. Sez-6 proteins affect dendritic arborization patterns and excitability of cortical pyramidal neurons. Neuron 56, 621–639 (2007).

34. Zhu, K., et al. Beta-Site Amyloid Precursor Protein Cleaving Enzyme 1 Inhibition Impairs Synaptic Plasticity via Seizure Protein 6. Biological psychiatry 83, 428–437 (2018).

35. Pigoni, M., et al. Seizure protein 6 and its homolog seizure 6-like protein are physiological substrates of BACE1 in neurons. Molecular neurodegeneration 11, 67 (2016).

36. Hidaka, C. & Mitsui, S. N-Glycosylation modulates filopodia-like protrusions induced by sez-6 through regulating the distribution of this protein on the cell surface. Biochemical and biophysical research communications 462, 346–351 (2015).

37. Gowen, A.M., et al. Role of microRNAs in the pathophysiology of addiction. Wiley interdisciplinary reviews. RNA, e1637 (2020).

38. Badiani, A., Belin, D., Epstein, D., Calu, D. & Shaham, Y. Opiate versus psychostimulant addiction: the differences do matter. Nature reviews. Neuroscience 12, 685–700 (2011).

39. Lenoir, M., Guillem, K., Koob, G.F. & Ahmed, S.H. Drug specificity in extended access cocaine and heroin self-administration. Addiction biology 17, 964–976 (2012).

40. John, B., et al. Human MicroRNA targets. PLoS biology 2, e363 (2004).

41. Sadakierska-Chudy, A., et al. Prolonged Induction of miR-212/132 and REST Expression in Rat Striatum Following Cocaine Self-Administration. Molecular neurobiology 54, 2241–2254 (2017).

42. Quinn, R.K., et al. Distinct miRNA expression in dorsal striatal subregions is associated with risk for addiction in rats. Translational psychiatry 5, e503 (2015).

43. Quinn, R.K., et al. Temporally specific miRNA expression patterns in the dorsal and ventral striatum of addiction-prone rats. Addiction biology 23, 631–642 (2018).

44. Shimizu-Nishikawa, K., Kajiwara, K., Kimura, M., Katsuki, M. & Sugaya, E. Cloning and expression of SEZ-6, a brain-specific and seizure-related cDNA. Brain research. Molecular brain research 28, 201–210 (1995).

45. Shimizu-Nishikawa, K., Kajiwara, K. & Sugaya, E. Cloning and characterization of seizure-related gene, SEZ-6. Biochemical and biophysical research communications 216, 382–389 (1995).

46. Herbst, R. & Nicklin, M.J. SEZ-6: promoter selectivity, genomic structure and localized expression in the brain. Brain research. Molecular brain research 44, 309–322 (1997).

47. Bahn, S., Volk, B. & Wisden, W. Kainate receptor gene expression in the developing rat brain. The Journal of neuroscience : the official journal of the Society for Neuroscience 14, 5525–5547 (1994).

48. Pigoni, M., et al. Seizure protein 6 controls glycosylation and trafficking of kainate receptor subunits GluK2 and GluK3. The EMBO journal 39, e103457 (2020).

49. Valbuena, S. & Lerma, J. Kainate Receptors, Homeostatic Gatekeepers of Synaptic Plasticity. Neuroscience 456, 17–26 (2021).

50. Delorme, R., et al. Frequency and transmission of glutamate receptors GRIK2 and GRIK3 polymorphisms in patients with obsessive compulsive disorder. Neuroreport 15, 699–702 (2004).

51. Xu, J., et al. Complete Disruption of the Kainate Receptor Gene Family Results in Corticostriatal Dysfunction in Mice. Cell reports 18, 1848–1857 (2017).

52. Lenoir, M., Augier, E., Vouillac, C. & Ahmed, S.H. A choice-based screening method for compulsive drug users in rats. Current protocols in neuroscience Chapter 9, Unit 9 44 (2013).

53. Lenoir, M. & Ahmed, S.H. Supply of a nondrug substitute reduces escalated heroin consumption. Neuropsychopharmacology : official publication of the American College of Neuropsychopharmacology 33, 2272–2282 (2008).

54. Mantsch, J.R., Yuferov, V., Mathieu-Kia, A.M., Ho, A. & Kreek, M.J. Effects of extended access to high versus low cocaine doses on self-administration, cocaine-induced reinstatement and brain mRNA levels in rats. Psychopharmacology 175, 26–36 (2004).

55. Livak, K.J. & Schmittgen, T.D. Analysis of relative gene expression data using real-time quantitative PCR and the 2(-Delta Delta C(T)) Method. Methods 25, 402–408 (2001).

56. Lenoir, M., Serre, F., Cantin, L. & Ahmed, S.H. Intense sweetness surpasses cocaine reward. PloS one 2, e698 (2007).

57. Paxinos, G. & Watson, C. The Rat Brain in Stereotaxic Coordinates: Compact 7th Edition. Academic Press 388 (2017).

58. Schroeder, A., et al. The RIN: an RNA integrity number for assigning integrity values to RNA measurements. BMC molecular biology 7, 3 (2006).

59. Agarwal, V., Bell, G.W., Nam, J.W. & Bartel, D.P. Predicting effective microRNA target sites in mammalian mRNAs. eLife 4 (2015).

60. Chen, Y. & Wang, X. miRDB: an online database for prediction of functional microRNA targets. Nucleic acids research 48, D127–D131 (2020).

61. Paraskevopoulou, M.D., et al. DIANA-microT web server v5.0: service integration into miRNA functional analysis workflows. Nucleic acids research 41, W169–173 (2013).

62. Vandesompele, J., et al. Accurate normalization of real-time quantitative RT-PCR data by geometric averaging of multiple internal control genes. Genome biology 3, RESEARCH0034 (2002).

63. Laemmli, U.K. Cleavage of structural proteins during the assembly of the head of bacteriophage T4. Nature 227, 680–685 (1970).

